# T-cell Receptor Diversity Estimates for Repertoires (TCRDivER) uses sequence similarity to find signatures of immune response

**DOI:** 10.1101/2021.01.11.417444

**Authors:** Milena Vujović, Paolo Marcatili, Benny Chain, Joseph Kaplinsky, Thomas Lars Andresen

## Abstract

We propose TCRDivER, a global approach to T-cell repertoire comparison using diversity profiles sensitive to both clone size and sequence similarity. As immunotherapies improve, the long standing biological interest in connecting outcome with T cell receptor (TCR) repertoire status has become more urgent. Here we show that new insights can be extracted from high throughput repertoire sequencing data. Most current efforts focus on identification of immunisation-specific sequence motifs or on monitoring changes in frequency of individual clones. Applying TCRDivER to murine spleen samples shows it characterises an additional dimension of repertoire variation, beyond conventional diversity estimates, allowing distinction between immunised and non-immunised samples. We further apply TCRDivER to repertoires from human blood. In both cases we show characteristic relationships between repertoire features. These reveal biologically interpretable relationships between sequence similarity and clonal expansions. We thereby demonstrate a new tool for investigation in clinical and research applications.

## Introduction

The T cell compartment of adaptive immunity plays a crucial role in cancer immunity, auto-immune and infectious diseases. Adaptive immune responses as a whole draw on diverse T-cell receptors. Due to the phenomenon of epitope spreading, T cells diversify their antigen-specific response by reacting to non-dominant epitopes present on the antigen, in addition to the main dominant epitope driven response ***Didona and Di Zenzo (2018); Vanderlugt and Miller (2002)***. On the other hand, it has been shown that T cells responding to the same epitope share more sequence similarity ***Dash et al. (2017)***. In addition, TCRs often exhibit cross-reactivity in order to ensure broad epitope recognition responses, despite the limited number of unique TCRs within each repertoire ***Petrova et al. (2012); Antunes et al. (2017); Bentzen and Hadrup (2019)***.

T cells are generated through the imprecise stochastic process of V(D)J recombination giving rise to 10^20^ or more possible TCR combinations ***Miles et al. (2011); Mora and Walczak (2018)***. The number of possible generated TCRs is much larger than those estimated to be present within any individual T cell repertoire ***Laydon et al. (2015); Robins et al. (2009, 2010)***. The complexity and composition of TCR repertoires makes it difficult to compare and stratify individuals based on immune status or to even establish a healthy baseline. T cells activate and proliferate upon antigen-specific contact, creating a complex mix of receptors. Initial experimental approaches, such as spectratyping ***Choi et al. (1989); Gorski et al. (1994); Memon et al. (2012); Ochsenreither et al. (2008)*** and flow cytometry ***Ciupe et al. (2013); Muraro et al. (2000)*** aimed to reveal oligoclonal expansions of T cells by tracking clonal sizes of CDR3s with the same length. However, they provided no insight into TCR similarity. Recent advances in high throughput sequencing (HTS) now allows characterisation of adaptive immune receptors in increasing depth and with improved quantitation. HTS methods supply information on both clonal sizes and sequence relatedness. However, this development has given rise to the need for summary measures to interpret the data generated by such experiments. Several methods have emerged to fulfil the demand to stratify repertoires either by the TCR antigen specificities ***Glanville et al. (2017); Dash et al. (2017); Sidhom et al. (2018)*** or by finding characteristics of TCR sequences ***Thomas et al. (2014a); Sun et al. (2017); Cinelli et al. (2017)***. Still, many of the employed methods aim to uncover epitope similarity without simultaneously examining T cell clonal expansions, or vice versa. Measures that capture the global repertoire structure by incorporating both characteristics of the adaptive immune response, could potentially be used to stratify patients for disease outcome or therapy. Thereby, transcending the notion of “public TCRs” into “public repertoire structures” responsible for therapeutic outcome.

A popular approach in characterisation of repertoires has been through measures of diversity. They have been widely used in evaluation of therapy and disease effects ***Twyman-Saint Victor et al. (2015); Rudqvist et al. (2018); Sherwood et al. (2013); Robert et al. (2014); Warren et al. (2011)*** or attempts at repertoire classification and diagnosis ***Carey et al. (2016); Chang et al. (2019); Robins et al. (2009)***. However, there are a variety of ways in which the intuitive idea of diversity can be formalised giving rise to ambiguity. In the naive sense, diversity is estimated based on the number of and clonal expansion of unique TCRs in a repertoire. Commonly used diversity estimates are richness (number of different TCR clones), clonality (number of expanded clones) and diversity indices such as Shannon entropy ***Spellerberg and Fedor (2003), Simpson SIMPSON (1949), Gini-Simpson Jost (2006)*** and Berger-Parker ***Berger and Parker (1970)*** index. Different diversity indices will weight expanded clones differently, thereby imposing a threshold on the clonal frequencies within the repertoire. Thus counting unique clones with species richness will give rare clones the same weight as expanded clones. Entropy, will give give more weight to expanded clones than to rare clones. No single index will capture all information about the clone size distribution. Notably, this ambiguity has led to no clear consensus which diversity index should be applied in practical cases of interpreting immune diversity ***Izraelson et al. (2018); Chiffelle et al. (2020)***. The approach of using individual diversity indices provides no repertoire characteristics truly independent of sample size ***Laydon et al. (2015)*** and can lead to erroneous conclusions on ordering repertoires (Figure 1 A.). Additionally, measures of diversity should not only rely on clone counts but should also account for sequence similarity of receptors.

**Figure 1.**
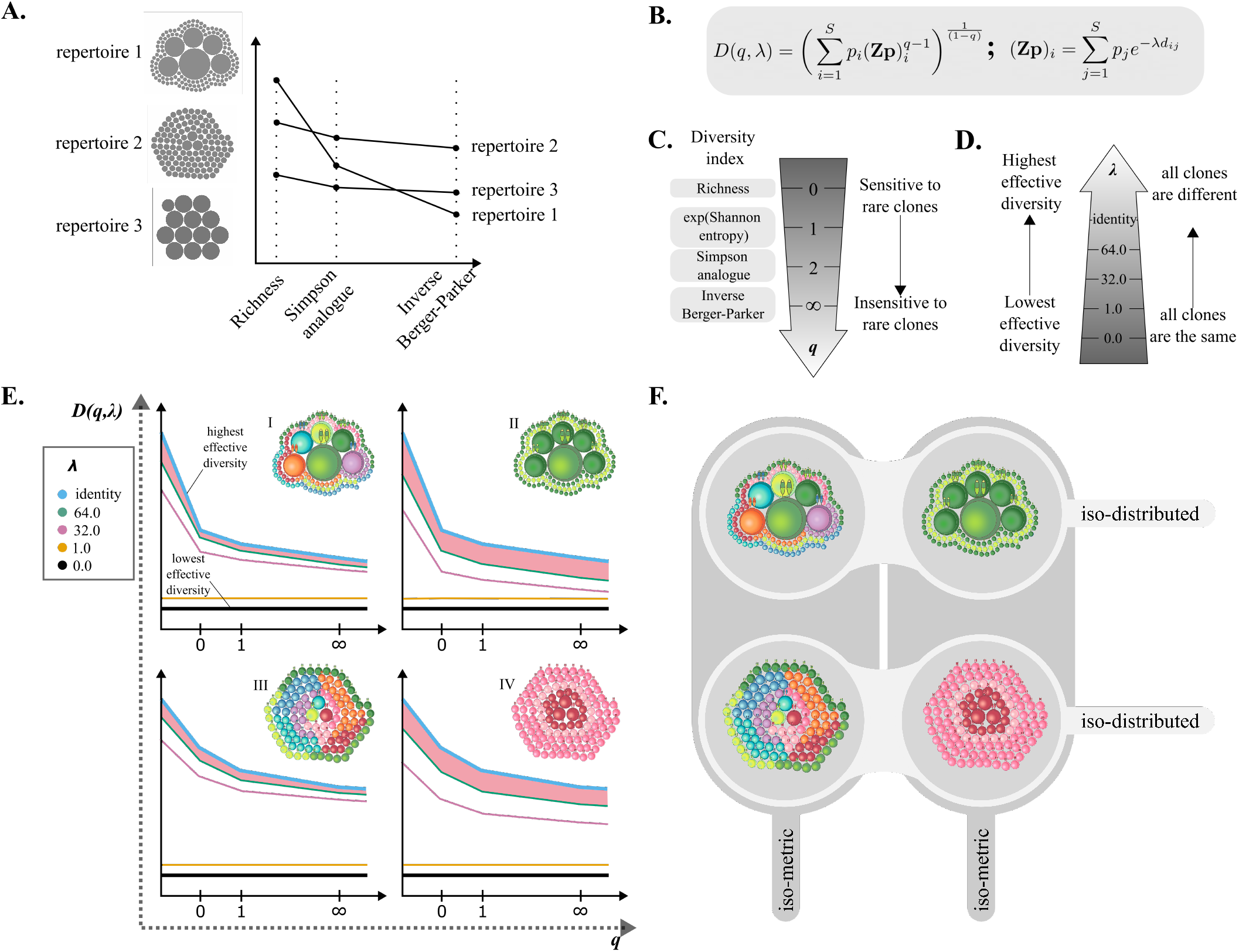
**A.** Naive diversity indices (richness, Simpson analogue and Inverse Berger-Parker) for three model repertoires. The repertoires contain the same total number of T cells, however they differ in the number of unique T cell clones and clone size distribution. Each T cell clone within the repertoire is presented as a grey circle and the size of each circle corresponds to its relative frequency. Repertoire 1 has the most uneven clonal distribution. Repertoires 2 and 3 have a more uniform distribution of clones. The number of unique clones falls from repertoire 1 through to repertoire 3. As different diversity indices are applied, the ordering of repertoires changes. **B.** Formula for calculating similarity scaled diversity *D*(*q, λ*). Here *p_i_* is the fraction of cells in clone *i*, **Z** is a similarity kernel between clones, *q* controls sensitivity to clone size, and *λ* controls sensitivity to sequence similarity. **C.** Relationship of some commonly used diversity indices to the naive diversity of order *q*. **D.** Effect of introducing the *λ* distance scaling into the diversity calculation. As *λ* increases the distance between clones increases until at *λ* = ∞ clones have no similarity i.e. the similarity kernel **Z** is the identity. **E.** Schematic representation of diversity profiles of four model repertoires are shown along the repertoires in the top right corner of each diversity profile. Repertoire composition is schematically represented with T cells of varying size and colour. Analogously to A. the size of individual cells corresponds to the clone frequency within the repertoire. More similar colouring indicates higher T cell sequence similarity in the repertoire (repertoires II and IV). Analogously, divergent colouring corresponds to higher sequence dissimilarity (repertoires I and III). **F.** Further explanation of structure differences between the four model repertoires in E. Repertoires sharing the same clone distributions are shown in rows (I = II and III = IV). Repertoires sharing same similarity relationships between T cell clones are shown in columns (I = III and II = IV).

The first problem of debatable usage of individual indices, can be surmounted by estimating them simultaneously in a single expression of diversity: the diversity of order *q*, *D*(*q*), which subsumes most of the commonly used indices ***Jost (2006, 2010)***. Accounting for the distribution of clone sizes, diversity can be estimated in the form of “diversity profiles” ***Greiff et al. (2015); Mora and Walczak (2016); Chiffelle et al. (2020)***. Such profiles define “effective numbers” of receptors when viewed at different resolutions, making use of a single parameter (*q*) to systematically shift focus from counting each unique clone to giving weight only to the largest clone in a repertoire (Figure 1 A, C). The use of diversity profiles gives insight into T cell clonal expansions, as the relationship between diversities calculated at different clonality weights *q* can be correlated to the ratio of common to rare clones ***Leinster and Cobbold (2012)***. This approach has been previously implemented for one B- and three T-cell repertoires in the work of ***Greiff et al. (2015)***. Keeping in mind that the study focused solely on clonal frequency, the authors report remarkable separation based on immunological status, in 3 out of 4 immune repertoire datasets. However, as naive diversity estimates only take clone frequencies into account they are not sensitive to minor polyclonal expansions of TCRs reacting to the same antigen which are mounting a unified front of reacting similar T cells.

The second problem, incorporating sequence similarity in diversity estimates, has been less thoroughly explored. One approach is to count clusters of similar receptors ***Sidhom et al. (2018)***. Another approach is to use an effective number with sensitivity to sequence similarity ***Arora et al. (2018)***. These approaches suffer from a similar limitation as use of a single diversity index in that they adopt either a single arbitrary cutoff or a single sequence similarity distance in their definition of effective number e.g. a single similarity corrected diversity index. Here we make use of approach used by ***Leinster and Cobbold (2012)*** in ecology to explore 2 dimensional profiles of effective numbers. Our approach of using similarity scaled diversity *D*(*q, λ*) allows for simultaneous characterisation of the clonal distributions and similarity of receptor repertoires (Figure 1 B). Instead of depending on a single parameter, *q*, our profiles depend on two parameters, *q* and *λ*. As in conventional diversity profiles, the *q* parameter probes the structure of the clone size distribution. The *λ* parameter plays an analogous role for sequence similarity (Figure 1 D.). As *λ* varies from infinity down to zero the effective diversity gradually merges together more and more similar sequences. Incorporating this additional aspect to the diversity estimation allows us not only to probe the clone size distribution, but also TCR similarity which may provide information on repertoire convergence through expansion of similar clones.

In this study, we showcase a new tool for estimating TCR repertoire diversity using similarity sensitive diversity estimates: TCRDivER. We apply TCRDivER to previously published murine TCR*β* sequence data from CD4^+^ T cells following immunization ***Sun et al. (2017)***. We show that TCRDivER, by simultaneously probing clonal expansion and sequence similarities reveals novel TCR repertoire traits. Using features of the similarity scaled diversity profile we detect differences in response to immunisation protocols at all sampling times, indicating unique features arise within repertoires can be detected as early as 5 days and persist for several months.

Notably we find strong nonlinear correlations between features of the similarity scaled diversity profiles, including feature which characterise the average distance between sequences or the balance between large and small clones. The strength of the correlation indicates biological constraints on repertoire development which couple together clone size with sequence similarity. The nonlinear shape of the correlations reveal that these features can exist in multiple discrete equilibria.

We validate our finding that TCR repertoires reside in a non-linear space on an independent dataset of human bulk TCR sequences extracted from non-small-cell lung cancer (NSCLC) patients following CTLA-4 blockade treatment ***Formenti et al. (2018)***. We show that by estimating repertoire diversity with TCRDivER we can unearth information which might allow us to understand more subtle differences between repertoires, stratify them and ultimately guide therapy regimes.

## Results

We analysed the frequency distribution and similarity of CDR_*β*_3 amino acid sequences in TCR repertoires previously published in ***Sun et al. (2017)***. Briefly, the dataset consists of CD4^+^ T-cell repertoires harvested from murine spleens following immunisation with Complete Freund’s Adjuvant (CFA) with or without the addition of Ovalbumin (OVA) antigen. The T cells were harvested post immunisation at three timepoints: early (days 5 and 14) and late (day 60). Additionally, we have analysed untreated mouse repertoires from the same study. We used TCRDivER to calculate diversities *D*(*q, λ*), with varying orders of *q* and *λ*. From these we constructed diversity profiles (divPs), which we present as graphs of the natural logarithm of diversity versus the varying order of q for each lambda. Key features were extracted for analysis as shown bellow.

To validate some of our findings we analysed a human TCR repertoire data set previously published in ***Formenti et al. (2018)***. In short, T cells were isolated from blood samples taken from non-small-cell lung cancer patients prior and post treatment with CTLA-4 blockade (ipilimumab) in combination with radiation therapy (RT). The obtained T cells were sequenced in bulk. We analysed these repertoires as with the murine data, using TCRDivER to construct diversity profiles for analysis.

### TCRDivER reveals unique TCR repertoire features

Examples of diversity profiles constructed for the murine samples are shown in figure 2. Each dataset was sampled for 50,000 sequences in order to eliminate effects of sequencing depth. For each sample a series of curves are plotted, corresponding to different values of *λ*. The constructed diversity profiles provide a graphically intuitive way to caputre the shape of a repertoire. Here we highlight some features of these plots in order to develop an understanding of how features of the plots map to structural and immunological characteristics of TCR repertoires.

**Figure 2.**
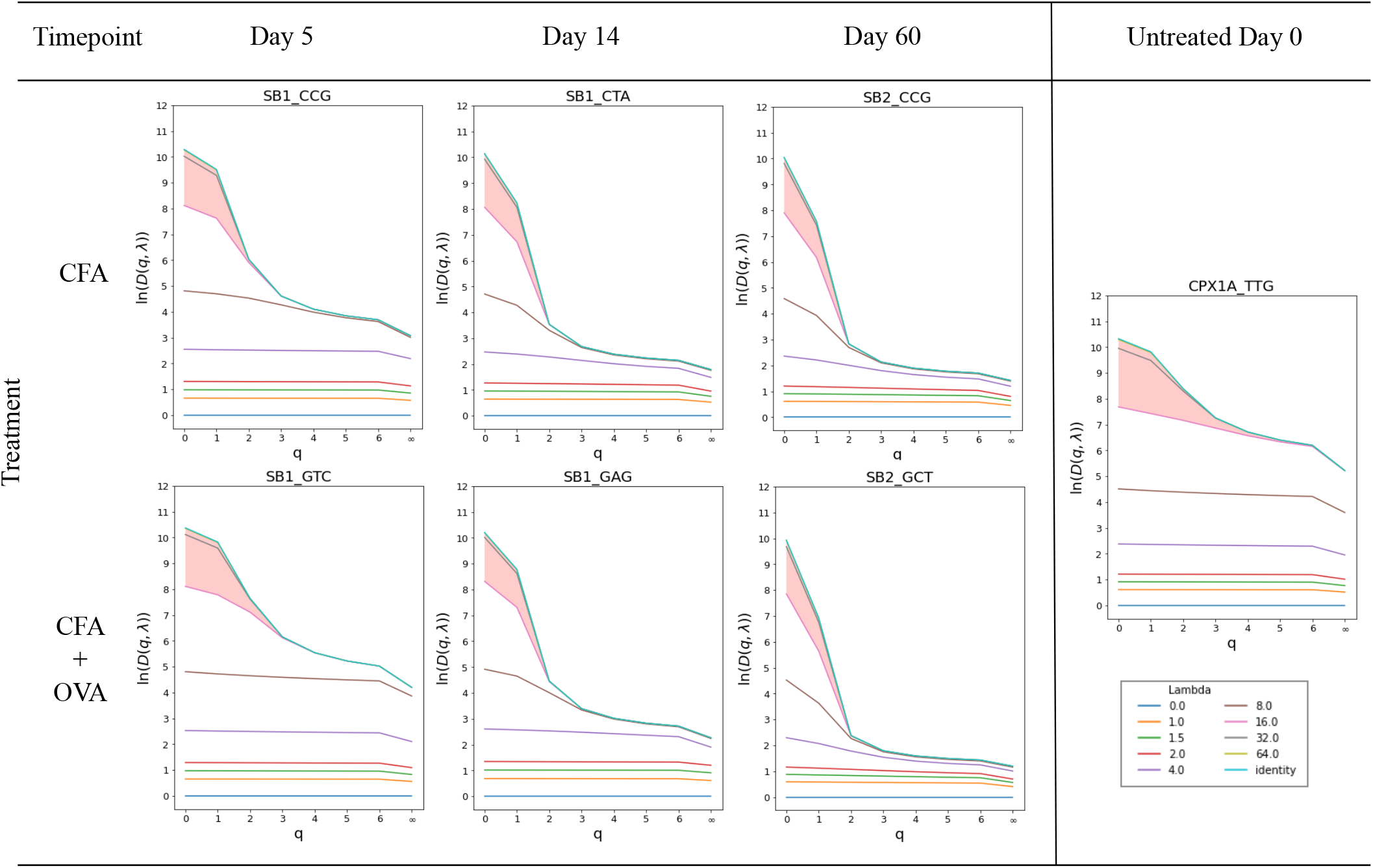
Diversity profiles (divPs) of calculated from CDR3 frequency within each repertoire and their similarity. DivPs of repertoires stemming from immunisation have been shown to the left, while the untreated is show on the right. Natural logarithm transformed values of diversity^*q*^ *D* for each calculated *λ* = 0.0, 1.0, 1.5, 2.0, 4.0, 8.0, 16.0, 32.0, 64.0 and identity versus the increasing order of *q*. The legend for all diversity profiles is shown at the bottom right. The highlighted area represents the area between *λ* 16.0 and identity curves. It highlights the change in repertoire CDR3 similarity unification for repertoires of different origin. Diversity profiles of only one sample per group are shown, the rest can be found in *Supplementary information - Section 1.1*.

In our framework the naive diversity profile corresponds to the case that the receptor of each T cell clone is considered totally distinct, with no consideration of similarity to other clones i.e. the highest effective diversity. In reality there will be some degree of functional overlap between clones, which will reduce the functional diversity below the naive value. The naive diversity (*λ* = ∞) is therefore a base case of maximal diversity. Atthe opposite extreme, *λ* = 0, all clones are considered considered functionally identical. Biologically, this would correspond to non-specific binding of TCRs to peptide-MHC complexes. In this case the functional diversity is therefore minimal and equals one. In each repertoire sample these two extreme cases can be seen bounding the profile (these curves are labelled in example figure 1 E.) The parameter *λ* interpolates between these two extreme cases, as the intermediate profiles in each sample correspond to intermediate values of *λ*.

We begin our account by highlighting features of the naive diversity, i.e. the upper bounding curves, in each sample. We plotted naive diversity profiles from all samples together showing that crossings in the range 0 ⩽ *q* ⩽ 2 are common events (See *Supplementary information - Section 1.2 and 1.6*). This confirmed in our data set that the ranking of repertoires based on a single value of *q* would indeed depend strongly on the chosen index (similar to what is shown in example figure 1 A.). We concluded that the previously mentioned justification for analysing profiles across a range of orders *q* is not merely theoretical.

The highest value of the naive diversity at *q* = 0 gives the number of unique TCR sequences observed in the sample of 50,000 sequences. At *q* = ∞ we read off the effective number of clones in the repertoire if it consisted only of the largest clones. The rate of fall of naive diversity as *q* rises therefore encodes information about the balance between larger and smaller clones. To characterise this we derived an expression forthegradient of the naive diversity at *q* = 1 and found that it is proportional to the variance of the clone size distribution i.e. the ratio of rare to common T cell clones in a repertoire (see Appendix 1 Evaluating the slope at *q* =1).

Notably, when examining the diversity profiles in Figure 2 and *Supplementary information - Section 1.1*, a sharper slope can be seen in the curves from repertoires that have been immunised compared to the untreated ones, especially for later time points. This is quantified in figure 3 A by plotting the slope between *q* = 0 and *q* =1 for each treatment group. The increasing value of the slope is indicative of an increased clonal expansion at later timepoints. The impact of clonal expansion in reducing diversity is seen to be mild at earlier time points after vaccination (5 and 14 days) and more marked at the day 60 time point.

**Figure 3.**
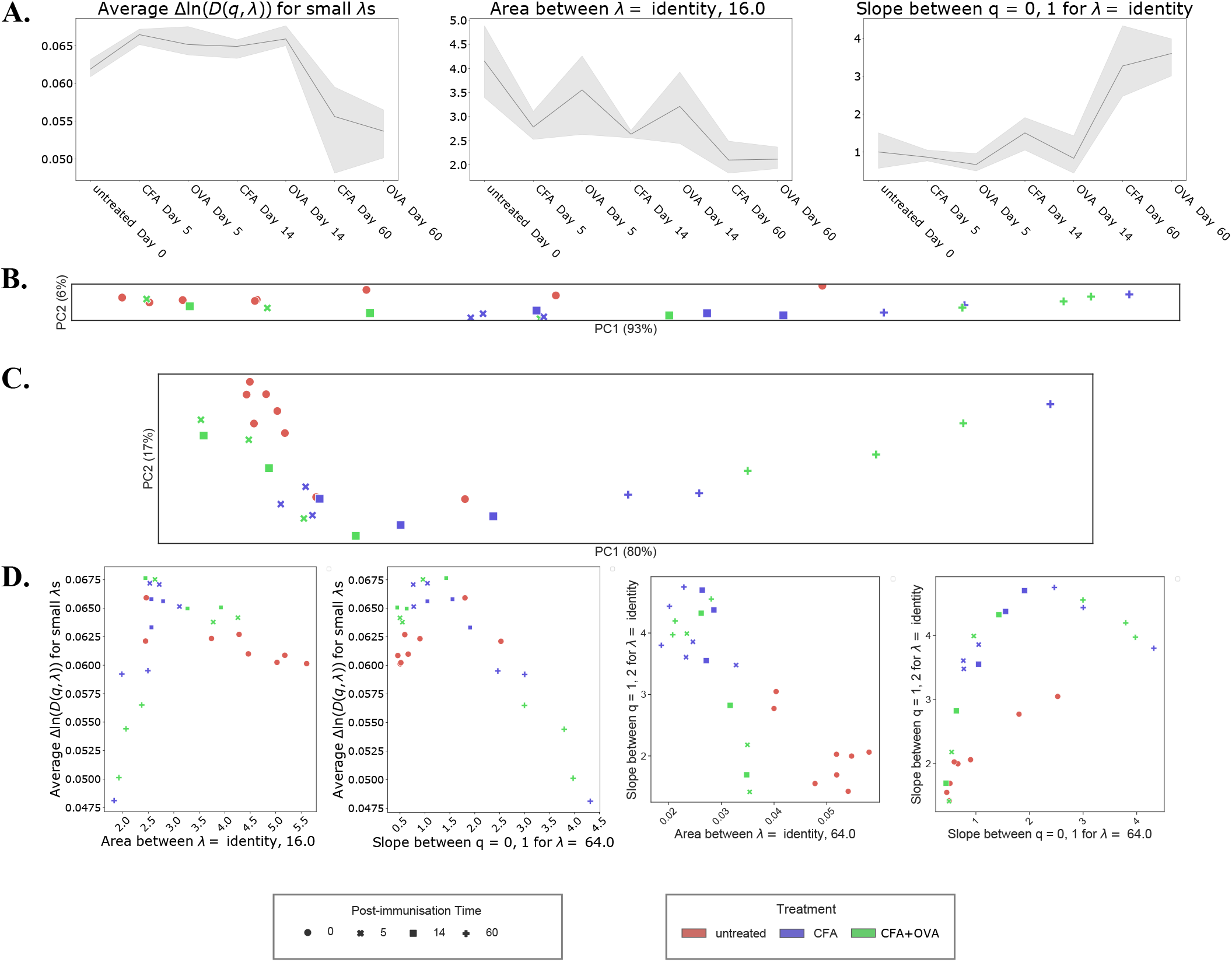
**A.** Trends of three features extracted from divPs are shown versus the treatment regime and timepoints ending with the latest timepoint. The features are, from left to right: average Δ ln (*D*(*q, λ*)) for small *λ*s, between curves of *λ* = identity and 16.0 and slope of *q* = 0 → 1 for value of *λ* identity. The line connects the mean values of the features for all samples within a group and the shaded area represents the confidence interval. **B.** PCA on naive diversity values^*q*^ *D*, i.e. *λ* = identity. The PCA plot aspect ratio has been adjusted and corresponds to variation explained by the first two principal components. **C.** PCA on features extracted from the diversity profiles constructed on the diversity values^*q*^ *D^Z^*. These include areas between all lambda curves, average Δln^*q*^ *D^Z^* for small *λ*s and slopes *q* = 0 → 1, *q* = 0 → 2 and *q* =1 → 2. As in C., the aspect ratio corresponds to variation found by PCA. **D.** Graphs showing relationships between some of the divP features. From left to right: average Δln(*D*(*q, λ*)) for small *λ*s is shown versus the area between curves of *λ* = identity and 16.0; average Δln(*D*(*q, λ*)) for small *λ*s is shown versus the slope of *q* = 0 → 1 for value of *λ* 64.0; slope of *q* = 1 → 2 for value of *λ* identity (i.e. naive diversity) is shown versus the area between curves of *λ* = identity and 64.0; slope of *q* = 1 → 2 for value of *λ* identity (i.e. naive diversity) is shown versus the slope of *q* = 0 → 1 for value of *λ* 64.0. The legend corresponds to figures B, C and D.

In order to explore the effects of similarity scaling on measures of diversity, we investigate in depth the features of similarity sensitive profiles with values of *λ* < ∞. A first feature of the similarity sensitive profiles occurs near the flat curve for *λ* = 0 at the bottom of the profile. We observed that for small values of *λ* the curves are approximately flat and are evenly spaced on a log scale. This phenomenon can be seen in figure 2 by observing that the spacing between the curves for *λ* = 0 (blue) and *λ* = 1 (orange) is equal to the spacing between the curves *λ* = 1 (orange) and *λ* = 2 (red). To further investigate the biological significance of this we derived an expression for the diversity at small *λ* using perturbation theory (See Appendix 2 Evaluation Δ*ln*(*D*(*q, λ*)) for small *λ*: Perturbation around *λ* = 0). Interestingly, this shows that the spacing is proportional to the mean distance between sequences in the repertoire. In particular, the spacing of profiles, which we denote Δ*ln*(*D*(*q, λ*)) at small *λ*, is dependent solely on the distance and frequency of CDR3s, and not on the weight *q*. Notably, this measure naturally integrates increases in similarity from both expansion of particular clones and selection of clones with similar sequence. We are therefore able to use these spacings to gain biological insight in to repertoire structure following immunisation. The spacing of profiles, Δ*ln*(*D*(*q, λ*)) at small *λ*, is presented for different treatment groups in figure 3 A. We concluded that while there may be a small rise in spacing at early time points after vaccination (5 and 14 days), there is a distinct decline of around 15% at day 60 indicating an increase of CDR3 similarity at later time points.

A second feature is the rate at which diversity falls as *λ* falls from *λ* = ∞ to lower values, where *λ* = ∞ characterises the naive profile. Unlike the case at small *λ*, the value of Δ*ln*(*D*(*q, λ*)) at large *λ* is no longer independent of *q*. Remembering that *λ* = ∞ corresponds to no effective clustering of similar sequences, large values of *λ* correspond to just a small amount of effective clustering counting together only the most similar clones. If such clustering produces a large fall in effective diversity then the repertoire must contain many similar clones. Conversely, if such clustering produces only a small fall in diversity then the clones must be spaced further apart.

Our measure is highlighted by the pink area in figure 2, defined as lying between the naive profile *λ* = ∞ and the profile for *λ* = 16. By using the area highlighted in the figure as the feature of interest we are effectively averaging over *q*. Notably, like Δ*ln*(*D*(*q, λ*)) at small *λ*, the area between *λ* = ∞ and *λ* = 16 is a probe of distance. However, in this case the weighting is toward similar clones. i.e. larger spacings correspond to more similarity of sequences in the repertoire. We have shown that, in the case of a uniformly distributed CDR3s in a repertoire, with the increase of similarity between CDR3s the area between the *λ* curves increases (see Appendix 2 Evaluation Δ*ln*(*D*(*q, λ*)) for larger *λ*s and it’s relationship to distance). In the case of natural repertoires the effect of clonal expansion will interplay with the similarity. The area between profiles for *λ* = ∞ and *λ* = 16, is presented for different treatment groups in figure 3 A. We concluded that there is a tendency to fall at the early time points after vaccination (5 and 14 days), with a further fall of comparable magnitude by day 60. As the area is influenced by *q* it is also closely connected to the slope of diversity curves and therefore clonal expansion. Therefore the fall of value of area cannot be attributed to a decrease in similarity at later time points. When other repertoire features are taken into account, such as the slope and the trend of Δ*ln*(*D*(*q, λ*)) at small *λ*, the decrease in area can be explained by the driving effect of clonal expansions at later time points.

Comparing the Δ*ln*(*D*(*q, λ*)) at small and large values of *λ* we were able to make some biological conclusions about the structure of the repertoires. At early time points there is a reduction in diversity of atypically similar (~ 1 amino acid difference) sequences. This may correspond to the expansion of responding clones with distinct sequences at the expense of background diversity. At early time points these expansions have little impact on the mean diversity, but by day 60 they have reduced mean diversity across the whole repertoire.

### TCRDivER features improve separation of biologically distinct repertoires

In order to test if the similarity scaled diversity profiles can be used to classify repertoires we used principal component analysis (PCA). We carried out this analysis first on the values for the naive diversity profiles alone and then for the all values in the complete similarity scaled diversity profile (Figure 3 B. and *Supplementary information - Section 1.3*.).

Both the naive and similarity scaled profiles show a strong PC1 which is driven by the expansion of large clones at the late day 60 time point. However, relative to the naive profile, PCA on the similarity scaled profiles shows more than twice the variance in PC2. In contrast to the naive profiles that give an effective 1 dimensional separation, the similarity scaled profiles are able to give a robust 2 dimensional separation. It can be seen that this allows for substantial separation of immunised vs. untreated controls in the second dimension (Figure 3 C.).

To validate the observation of improved PCA separation, we analysed an additional data set of human TCR repertoires. Briefly, this arose from blood samples collected prior and post immunotherapy from 40 patients diagnosed with stage 4 non-small-cell lung carcinoma (NSCLC). The therapy consisted of a regime of radiation and administering CTLA-4 blockade. After therapy completion each patient was categorised according to RECIST response criteria into four categories based on the therapy outcome (for further details see Section Data Acquisition and Description). We analysed the data as before by calculating the diversity *D*(*q, λ*) and constructing diversity profiles (see *Supplementary information - Section 2.1*.).

To test if the features we have identified are useful in capturing key dimensions of variation in the similarity scaled diversity profiles we then extracted these features and carried out PCA on the features rather the raw values. These include all the areas between different *λ* curves, average values of Δ ln (*D*(*q, λ*)) for small *λ*s and slopes for *q* = 0 → 1, *q* = 1 → 2 and *q* = 0 → 2. To further mitigate the effect of repertoire size on the analysis we have log-transformed the values of *D*(*q, λ*) prior to feature extraction. The analysed features are therefore ratios and relationships of the natural logarithm of *D*(*q, λ*). The results of the PCA analysis on the human can be seen in *Supplementary information - Section 2.2*.

In both the mouse and human data sets we found that the PCA on features was qualitatively similar to that on the full diversity profile. However, the variance explained by PC1 was reduced while that explained by PC2 was increased, leading to improved 2 dimensional separations. In the case of the mouse data the effect was modest, while in the human data it was more substantial, leading to an increase in variance explained by PC2 from 11% to 20%.

The reduction of variance explained in PC1 indicates that these features are indeed acting as useful summaries of redundant (linearly correlated) information in the full profile. Therefore making use of such features, rather than the raw profile values may help reduce experimental noise and improve robustness.

### Non-linear relationships between TCRDivER features are driven by the structure of repertoires

Because a significant proportion of the variance is not captured by PC1 alone, there must be some non-linear relationships between similarity sensitive diversity profile features. We therefore decided to look more closely at pairwise relationships between these features, as shown in figure 3 D.

This revealed two interesting characteristics of the relationships. Firstly, while many of the plots are clearly non-linear, they lie on surprisingly tight curves. This indicates that there is some constraint at work in the structure of the repertoire meaning that the value of one feature tightly constrains the value of other features. However, the second interesting characteristic is that these constraints are not unique.

An example is the relationship between Δ*ln*(*D*(*q, λ*)) at small *λ* and the area between profiles for *λ* = ∞ and *λ* = 16, as shown in the first panel of figure 3 D. As noted above, these both depend on the distances between sequences in the repertoire. Rising Δ*ln*(*D*(*q, λ*)) at small *λ* indicates mean distances between sequences in the repertoire increasing. Falling values of the area indicate fewer clones at very close distances to another clone. In a simple ideal case where the clone size and distance distributions are uniform these two effects perfectly coincide. We illustrate this using model data in Appendix 2 Evaluation Δ*ln*(*D*(*q, λ*)) for larger *λ*s and it’s relationship to distance. In real data with non-uniform clone size and distance distributions *D*(*q, λ*) provides a measure where the differential effect of changes in the most similar sequences and changes in comparisons across the repertoire as a whole can be characterised.

For data from the mouse immunisation experiments, figure 3 D illustrates a non-linear change point in the relationship between Δ*ln*(*D*(*q, λ*)) at small *λ* and area the between profiles for *λ* = ∞ and *λ* = 16. Above a threshold in area for larger *λ*s of around 2.5 we see the expected behaviour for the simple ideal case. Below this threshold we see the more complex phenomenon where a smaller mean distance between sequences goes with fewer small inter-clone distances. This is most developed in the samples from late time points where expanded clones are present. Within these clones the distances between sequences will be zero, pushing down the mean distance between sequences. At the same time the presence of large expanded clones means that these clones are less likely to have close neighbours.

The example of the relationship between Δ*ln*(*D*(*q, λ*)) at small *λ* and the area between profiles for *λ* = ∞ and *λ* = 16 illustrates a non-unique constraint. A value of 0.06 for Δ*ln*(*D*(*q, λ*)) at small *λ* constrains the value of the area between profiles for *λ* = ∞ and *λ* = 16 but to one of two possible values - either around 2.0 or around 5.5. This again emphasises the importance of taking multiple features of the diversity profile to more fully characterise repertoire structure. Similar non-unique constraints can be seen in the other panels of figure 3 D.

The mathematical form of the diversity we have adopted does impose some restrictions on possible diversity profiles. For example, the effective diversity must always fall (or stay constant) as *q* rises, reflecting the down weighting of small clones. To test if these relationships might be an artefact imposed by the mathematical form of the diversity we replaced the clone size distribution with pseudo-random numbers while keeping the distance matrix fixed, with results shown in figure 4. This eliminated observed correlations, showing that the correlations are not mathematically necessary. To confirm that the correlations are not a product of the particular distance definition adopted we repeated the analysis using an alternative metric based on amino acid properties. This shows qualitatively similar correlations (See *Supplementary information - Section 1.9*).

**Figure 4.**
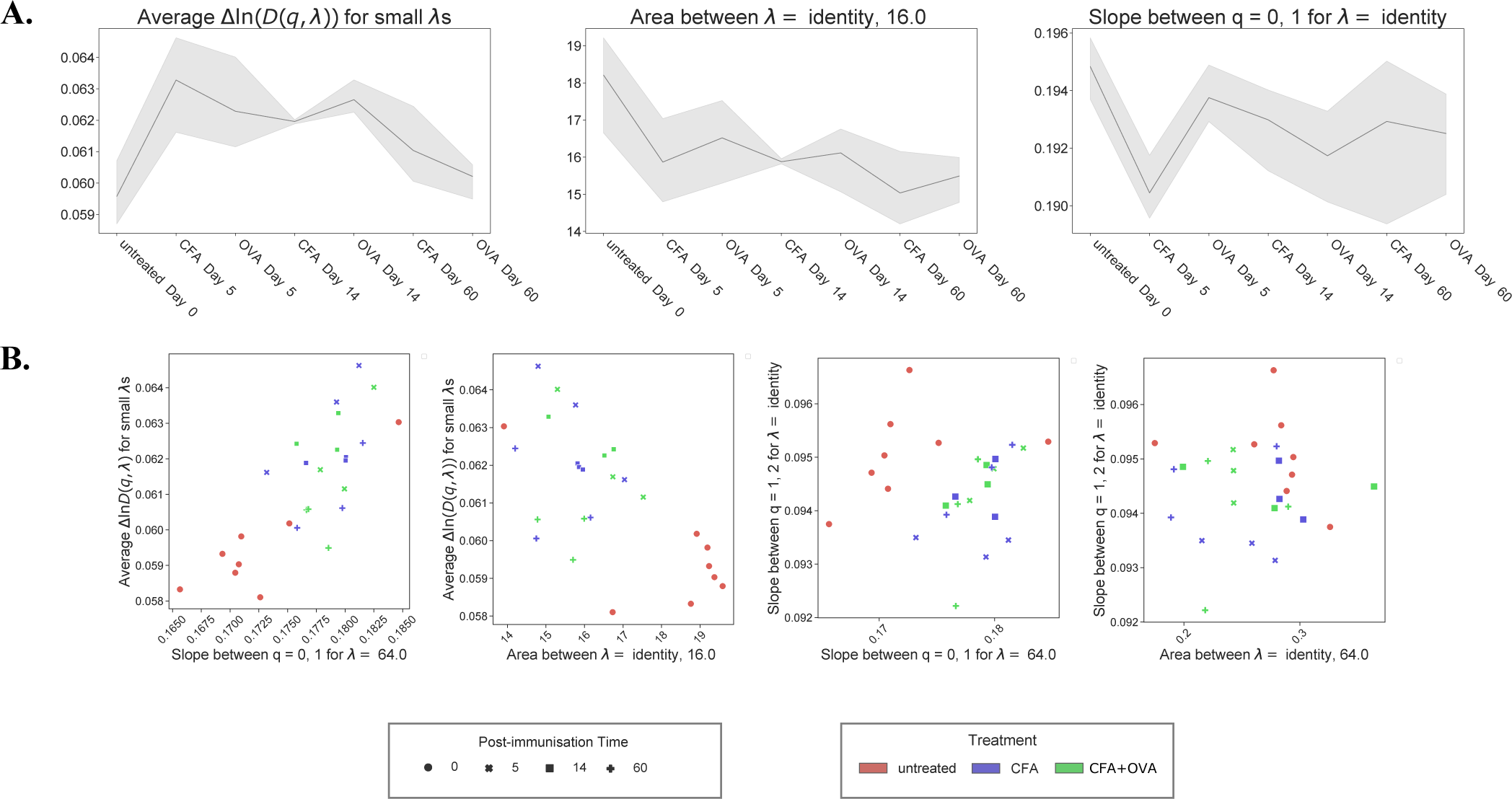
**A.** Trends of three features for the randomised murine dataset are shown versus the treatment regime and timepoints ending with the latest timepoint. The features are, from left to right: average Δln(*D*(*q, λ*)) for small *λ*s, between curves of *λ* = identity and 16.0 and slope of *q* = 0 → 1 for value of *λ* identity. The line connects the mean values of the features for all samples within a group and the shaded area represents the confidence interval. **B.** Graphs showing relationships between some of the divP features of the murine dataset with random frequencies. From left to right: average Δln (*D*(*q, λ*)) for small *λ*s is shown versus the slope of *q* = 0 → 1 for value of *λ* 64.0; average Δln(*D*(*q, λ*)) for small *λ*s is shown versus the area between curves of *λ* = identity and 16.0; slope of *q* = 1 → 2 for value of *λ* identity (i.e. naive diversity) is shown versus the slope of *q* = 0 → 1 for value of *λ* 64.0; slope of *q* = 1 → 2 for value of *λ* identity (i.e. naive diversity) is shown versus the area between curves of *λ* = identity and 64.0.

Turning to the human data set we found that pairwise relationships between features were quantitatively quite different that the mouse data, but displayed the same characteristics of lying on curves and giving rise to non-unique constraints (Figure 5 A.). Given differences in species, tissue and treatment, it is unsurprising that the range of repertoire structures observed differs considerably. At least some of these differences are captured in features of the similarity scaled diversity profiles. Despite these differences, the human data corroborates the notion that regardless of the immunisation strategy and dataset (human or murine), the natural TCR repertoires reside in a subspace governed by a complex interplay of TCR clonality and similarity.

**Figure 5.**
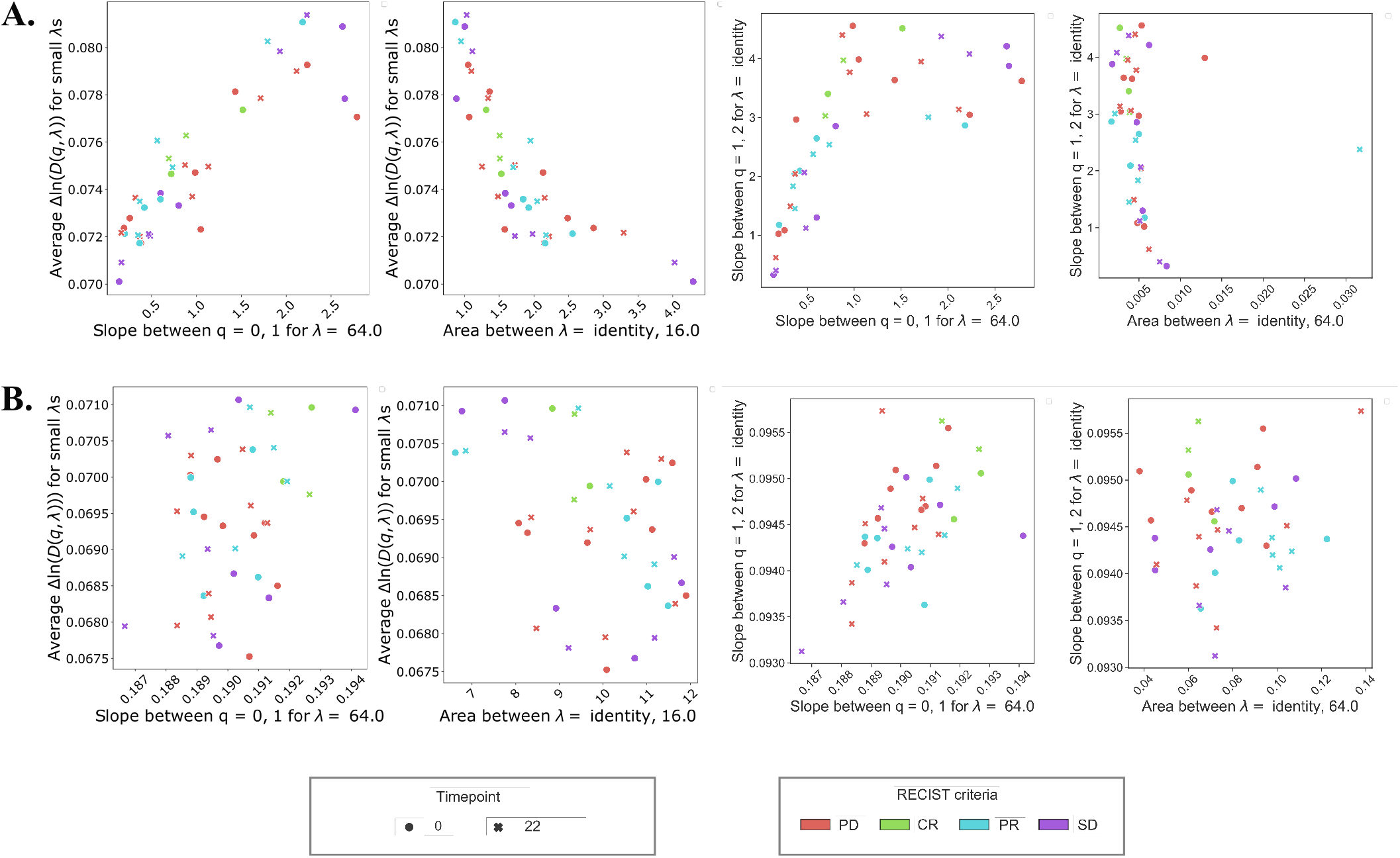
**A.** Graphs showing relationships between some of the divP features extracted from diPs of the human dataset. From left to right: average Δ ln (*D*(*q, λ*)) for small *λ*s is shown versus the slope of *q* = 0 → 1 for value of *λ* 64.0; average Δ ln (*D*(*q, λ*)) for small *λ*s is shown versus the area between curves of *λ* = identity and 16.0; slope of *q* = 1 → 2 for value of *λ* identity (i.e. naive diversity) is shown versus the slope of *q* = 0 → 1 for value of *λ* 64.0; slope of *q* = 1 → 2 for value of *λ* identity (i.e. naive diversity) is shown versus the area between curves of *λ* = identity and 64.0. **B.** As A., but comparing features from the human dataset with randomized frequencies. The legend corresponds to both A. and B.

## Discussion

The complex structure of immune repertoires makes them challenging to compare and classify. Previous work making use of sequence information to understand TCR repertoires has focused on determining the antigen specificity of particular sequences. In contrast, TCRDivER is able to make use of sequence information to reveal structural similarity between repertoires that have little or no sequence overlap. With 2 tunable parameters TCRDivER becomes an effective computational microscope, able to focus on different scales of structure in the immune repertoire. The resulting diversity profiles then provide highly interpretable summaries of global multiscale structure.

Here we provide a proof-of-concept study in application of similarity scaled diversity estimates to TCR repertoire analysis. We have applied principal components analysis as a means of qualitative repertoire stratification and shown it is able to capture variation in the diversity profiles which characterise immunisation history. The similarity scaled diversity profiles are themselves quite rich summaries and in the future we anticipate that they may be subject to more sophisticated machine learning techniques. At present this possibility is limited by the number of available samples for which comparable data is available. The need to define a similarity scaled diversity index with a parameter *λ* arises naturally from a desire to generalise the idea of naive diversity profiles. While there may be a variety of ways to define such a generalisation, it should be noted that the specific form of equation 7 (also shown in figure 1 B.) that we use possesses important abstract mathematical properties that will enable further investigation ***Tom Leinster (2013)***.

As has previously been shown ***Greiff et al. (2015)*** the use of a single diversity index to rank or classify repertoires is not robust, since the ranking will depend on the index selected. The use of naive diversity profiles is a step forward in so far as they reflect the contribution of both large and small clones. However, as applied to real TCR data naive diversity profiles typically give a single effective dimension of separation. This is reflected in our results that show 93% of variance explained by PC1 carried out on the naive diversity profile. The common choice to model clone size distributions using power laws ***Altan-Bonnet et al. (2020)*** that have a single tunable parameter further supports the idea that range of possible biological distributions is essentially 1 dimensional. In this case, any practical classification of repertoires based on naive diversity will be based on a 1 dimensional separation.

The novel features identified in our similarity scaled diversity profiles provide a genuine second dimension of variation in the structure of the repertoire based on sequence similarity. As shown in our PCA analysis this opens the practical possibility of 2 dimensional separations of repertoires which are inherently more powerful. Notably, this approach can make use of sequence data to classify repertoires together even when they share no similar sequences.

The striking relationships between similarity profile features appear to reflect biological structure in the TCR repertoires. We hypothesise that the non-linear relationships we observe reflect the range of possible biological variation and are thus analogous to the way in which clone size distributions are well approximated by power laws. Our observation motivates investigations of extending of power law distributions to include description of sequence similarity.

Any calculation involving an all-against-all comparison will inevitably scale with the square of the number sequences. Because TCRDivER is parallelisable, with the 50,000 sequences per sample analysed here it is very practical to run on commonly available computer clusters. When distributed over 8 cores of an Intel Xeon Gold 6126 2.60 GHz processor, each repertoire computation took under 6 hours. There is scope for several-fold speed up, including optimisation of the distance function. A possible gain would come from replacement of the exact all-against-all comparison at high lambda with comparison against approximate k-nearest neighbours using a ball tree algorithm. However, at low lambda values the all-against-all comparison cannot be avoided reflecting the way in which the similarity scaled diversity incorporates genuinely global information about the repertoire.

The form of equation 7 is motivated by rather general mathematical considerations, but these still leave the metric used to compare sequences undetermined. There is no ‘true’ metric, in the sense that a assigning a single number to the distance between two sequences cannot fully capture all the ways in which binding affinities vary. In our work we used two metrics which plausibly reflect biological functional similarity (through use of evolutionary data in BLOSUM45 matrix) and biochemical similarity (thorough the Atchley factors). While these gave qualitatively similar results, the best metric to use for a given question is an open question in the field of TCR analysis as a whole.

While in this study we have applied similarity scaled diversity profiles to TCR repertoires, we believe that the same concept should also be applicable for understanding antibody repertoires. The features of similarity scaled diversity profiles can easily be translated in to properties of the repertoire. The functional biological significance of the similarity scaled diversity (as indeed the naive diversity) is likely to be more variable and subject to experimental investigation. TCRDivER provides an important tool to enable those investigations.

## Materials and methods

### Data Acquisition and Description

The murine dataset consits of previously published data that has been analysed as part of a larger dataset in the work by ***Sun et al. (2017)***. Briefly, CD4^+^ T cells were isolated from spleens of 18 C57BL/6 mice immunised with Complete Freund’s adjuvant (CFA) with or without an addition of Ovalbumin antigen (OVA). The samples were collected at different times post-immunisation: at day 5 and 14 (early timepoints) and day 60 (late timepoint). In addition, CD4^+^ T cells were collected from 8 healthy unimmunised mice prior to study start. An overview of the dataset is given in Table 1. We have received the dataset already analysed with Decombinator, which described in depth in ***Thomas et al. (2013, 2014b)***. The data we have analysed consisted of a list of CDR3 sequences present in each sample. The raw fastq files are available at http://www.ncbi.nlm.nih.gov/sra/?term=SRP075893.

**Table 1.** Murine Dataset overview.

source data 1. Overview of samples available in the analysed dataset, a part of a previously published dataset by *Sun et al.
| Treatment | Sample collection time (days) | Number of mice |
| --- | --- | --- |
| CFA | 5 | 3 |
| CFA | 14 | 3 |
| CFA | 60 | 3 |
| CFA+OVA | 5 | 3 |
| CFA+OVA | 14 | 3 |
| CFA+OVA | 60 | 3 |
| Non-immunised | 0 | 8 |

Additionally, we have analysed a human TCR dataset previously analysed by ***Formenti et al. (2018)***. The participants of the study were 39 patients diagnosed with metastatic non-small-cell lung cancer (NSCLC). They were treated with daily radiation therapy regimen in two phases of the trial (phase I - 6Gy × 5 and phase II 9.5 Gy ×)and intravenous ipilimumab(CTLA-4 blockade) following the first radiation treatment and subsequently repeated every 3 weeks for four cycles. The assessment of patient treatment response was performed with PET/CT scans at day 88 and evaluated using Response Criteria In Solid Tumors (RECIST). The patients were then classified, according to RECIST, into complete responders (CR), partial responders (PR) with tumour decrease in size ⩽ 30%, stable disease (SD) with insufficient shrinkage to qualify for any of the other criteria, and progressive disease (PD) with increase in size > 20% or appearance of new lesions. Out of 39 patients 20 were evaluable at day 88. Serial blood samples for peripheral blood mononuclear cells (PBMCs) were collected at baseline (day 0), and on days 22, 43, 64, and 88. The isolated PBMC were subjected to amplification and sequencing of bulk TCRj*β* CDR3 regions by Adaptive Biotechnologies. We have obtained the data from the Adaptive Bioctechnologies ImmunoSEQ database ***Imm (????)***. Since the samples collected at later timepoints (day 43 and onward) were not available for all of the 20 evaluable patients, we have restricted our analysis to samples collected at baseline and day 22 of treatment. An overview of samples included in our analysis is given in Table 2.

**Table 2.** Human Dataset overview.

| RECIST criteria | Sample collection time (days) | Number of patients |
| --- | --- | --- |
| PD | 0 | 8 |
| PD | 22 | 8 |
| SD | 0 | 5 |
| SD | 22 | 5 |
| PR | 0 | 5 |
| PR | 22 | 5 |
| CR | 0 | 2 |
| CR | 22 | 2 |

Overview of samples used from the dataset previously published by *Formenti et al. (2018)*. In total 40 human PMBC samples were sequenced for TCR $\beta$ CDR3. Sample collection time has been given as the number of days after start of therapy, with 0 being baseline prior to first treatment.

### Subsampling to reduce computational load

We have only considered “In” frame reads of CDR3s in our analysis. In order to reduce the computational load of calculating pairwise similarity between a large number of CDR3 regions (order of magnitude ≈ 10^5^), we have performed subsampling prior to analysis. We will refer to individual CDR3 sequences as sequences, and a collection of identical CDR3 sequences as a clone. Each repertoire was down sampled to contain 50000 CDR3 sequences. The sampling was random, based on the original CDR3 frequency distribution. After sampling the number of CDR3 clones was ⩽ 50000. An overview of number of unique clones forthe murine and human dataset is given in Table 3 and 4, respectively. For each sample, the CDR3 clone count (number of identical CDR3s within a clone) was transformed into frequencies, so that the all the CDR3 clone frequencies sum up to 1:

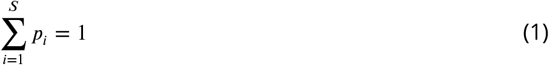

where *p_i_* is the frequency of the *i*-eth CDR3 clone in the repertoire. The clone CDR3 amino acid sequences with their respective frequencies were used as input further downstream.

**Table 3.**
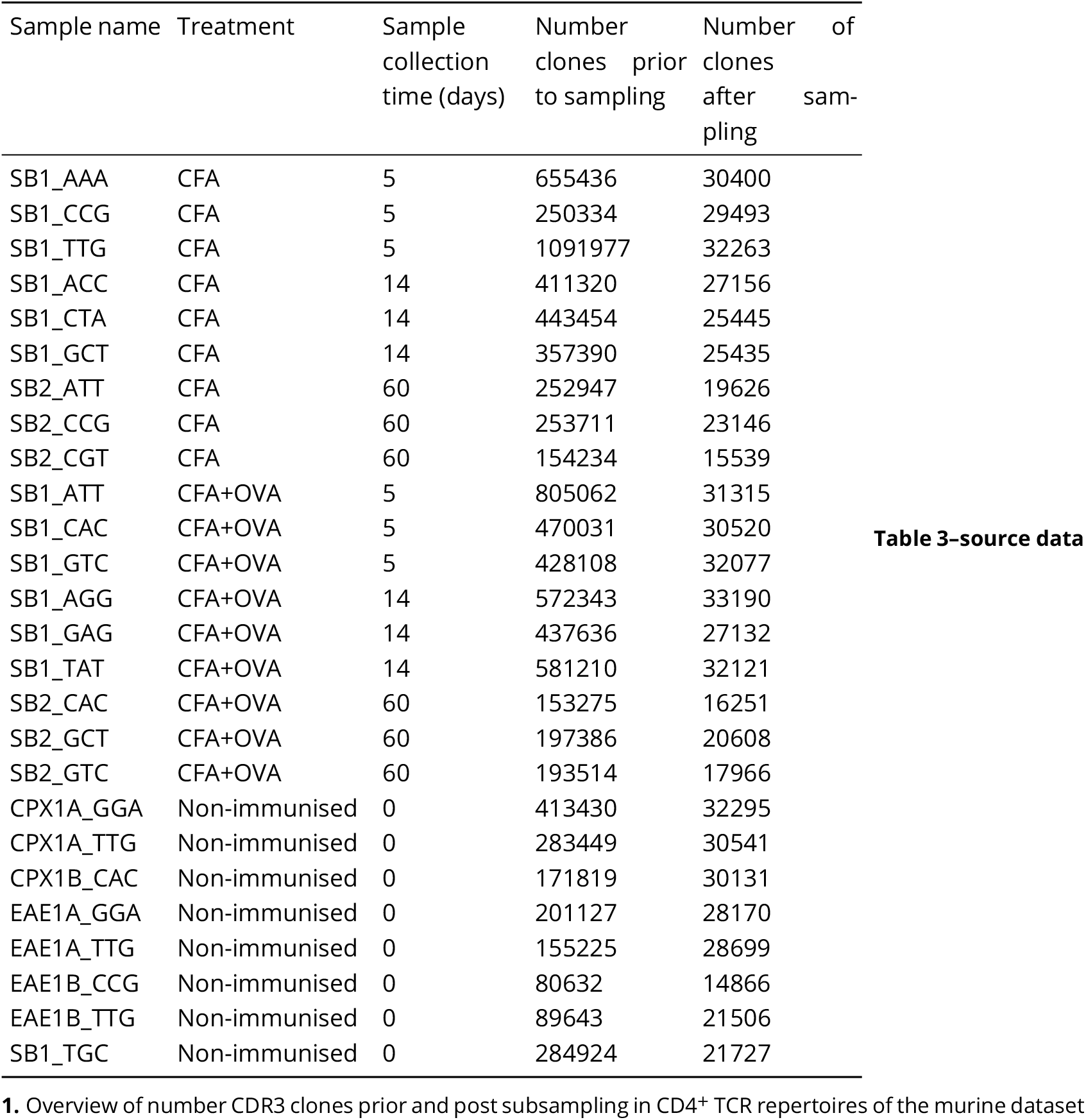
Murine Dataset Subsampling.

**Table 4.**
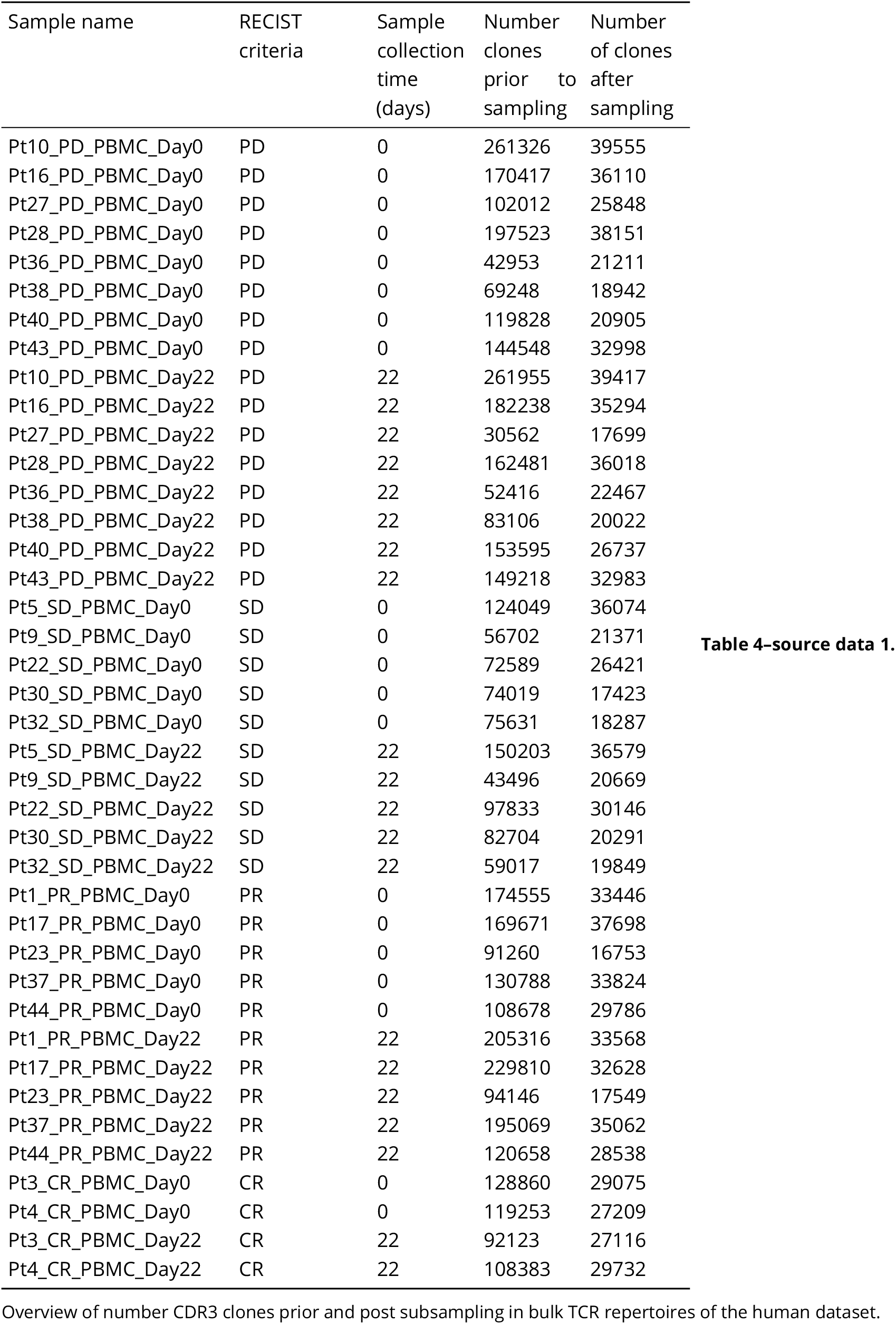
Human Dataset Subsampling.

### Calculating the Distance Matrix

In order to reduce computational time and memory usage of calculating pairwise comparison, we have divided the distance matrix into smaller portions (“chunks”) which are separately calculated. The algorithm takes in the list of *S* CDR3 clones, and splits it up into *n* sub lists of predefined length, default value is 100. For each sublist pairwise comparisons are made between the CDR3s in the sublist versus all the CDR3s in the original list. An example of the distance matrix calculation can be seen in Figure 6. Global alignment between two CDR3 clone sequences was performed with a gap penalty of 10 and scored using the BLOSUM45 ***Henikoff and Henikoff (1992)*** substitution matrix. The alignments were created using the PairwiseAligner function in the Bio.Align package within Python3 ***Van Rossum and Drake (2009)***. The distance between two CDR3s, *d*(*CDR3_i_, CDR3_j_*), was calculated based on the alignment scores:

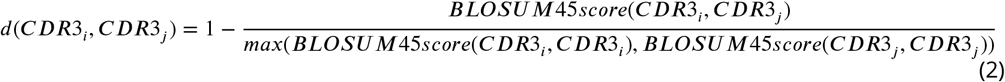

**Figure 6.**
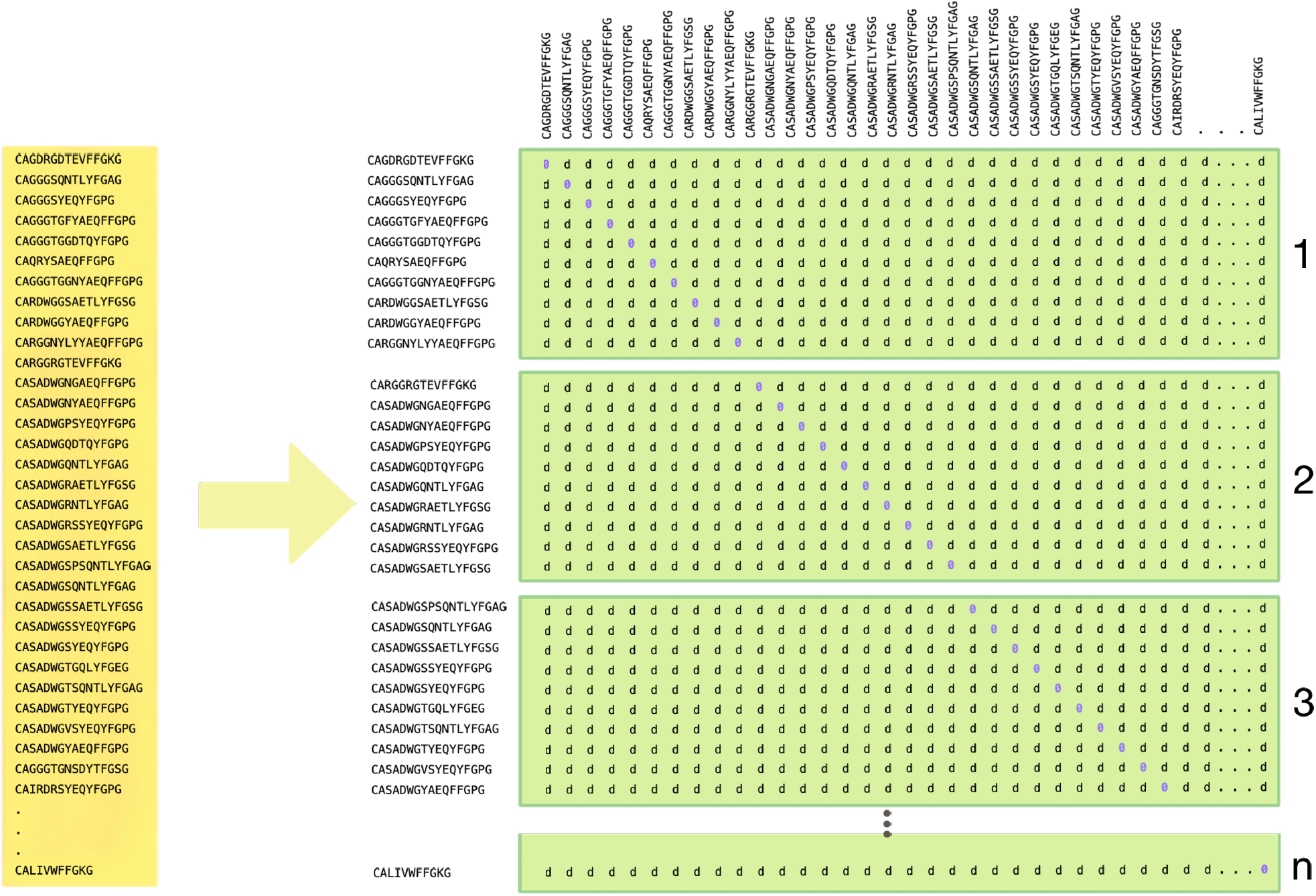
An overview of the calculation of the distance matrix. The list of CDR3 sequences is divided into lists of equal length, here 10 sequences, the default value is 100. These 10 CDR3s are then pairwise compared with all the other CDR3s in the total list of CDR3 sequences. Each portion of the distance matrix i.e. chunk has 10 rows and S columns, where S corresponds to the total number of CDR3 clone sequences. In the end there are n chunks, where n is equal to the floored division of total number of sequences by the length of chunk 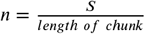. Distances *d* are *d*(*CDR3_i_, CDR3_j_*) calculated based on the BLOSUM45 alignment score. The diagonal of the combined distance matrix is 0.

Alternatively, we have also employed a biochemical scoring based on the Atchley factors ***Atchley et al. (2005)***. The factors are based on biochemical properties of amino acid residues that are grouped and transformed into five Atchley factors. For each CDR3 an average value for individual Atchley factors was computed. The distance between two CDR3 sequences is then calculated as an Euclidean distance between the averaged five Atchley factors.

The obtained distance matrix is mathematically connected to the similarity kernel Z as shown in equation 3

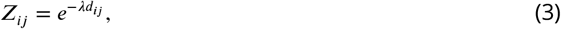

where the *λ* provides scalling.

### Calculating Naive Diversity

Naive diversity does not take similarity into account and is calculated based on the frequencies of CDR3 clones within the repertoire ***Jost (2006, 2010)***:

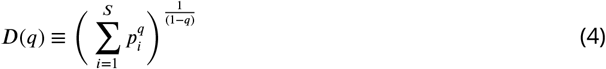

where *p_i_* is the frequency of the *i*-eth CDR3 clone and *q* is the diversity order, as diversity indices are a functions of 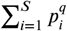. We have chosen a list of *q*s which subsume the use of some common diversity indices such as 0,1,2 and ∞, which correspond to the richness, exponent of the Shannon diversity ***Spellerberg and Fedor (2003)***, Simpson ***SIMPSON (1949); Jost (2006)*** and Berger-Parker ***Berger and Parker (1970)*** index, respectively. To extend our surveying the clone size distribution space we have also added 3, 4, 5 and 6^th^ order of diversity. For values of *q* = {1, ∞} the diversity was calculated as the limit of *q* approaching the values of 0 and ∞.

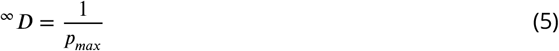

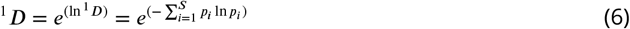

A detailed derivation of the equations is provided in Appendix 1.

### Calculating Similarity scaled Diversity

Similarity scaled diversity of order *q* takes CDR3 clone sequence distances *d*(*CDR3_i_, CDR3_j_*) along with their respective frequencies. We have adapted the method of calculating similarity-sensitive diversity measures, as proposed by ***Leinster and Cobbold (2012)***. We have again chosen a list of *q*s, *q* = 0, 1, 2, 3, 4, 5, 6, ∞, in the same manner as when calculating the naive diversity. Furthering the method, ***Leinster and Cobbold (2012)***, propose the use of similarity-sensitive diversity measures^*q*^ *D^Z^* (Equation 7), dependent on relative abundances and species similarity data as distance *d_ij_*. We introduced an alteration of the original approach is introducing the similarity scaling factor *λ*, which allows us to weight TCR distances in much the same way as we do clone sizes (Figure 1 D.). Here we choose a list of values *λ* = {0.0, 0.1, 0.2, 0.25, 0.3, 0.4, 0.5, 0.75, 1.0, 1.5, 2.0, 4.0, 8.0, 16.0, 32.0, 64.0 }. The value of *λ* = identity corresponds to the naive diversity calculation, *D*(*q*).

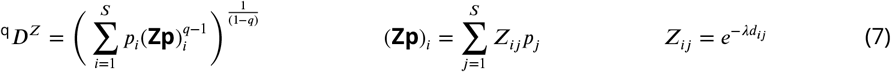

### Algorithm overview

An overall scheme of the TCRDivER algorithm is given in Figure 7. The algorithm has been implemented within Python (v3.6) ***Van Rossum and Drake (2009)*** and it’s freely available at https://github.com/sciencisto/TCRDivER.

**Figure 7.**
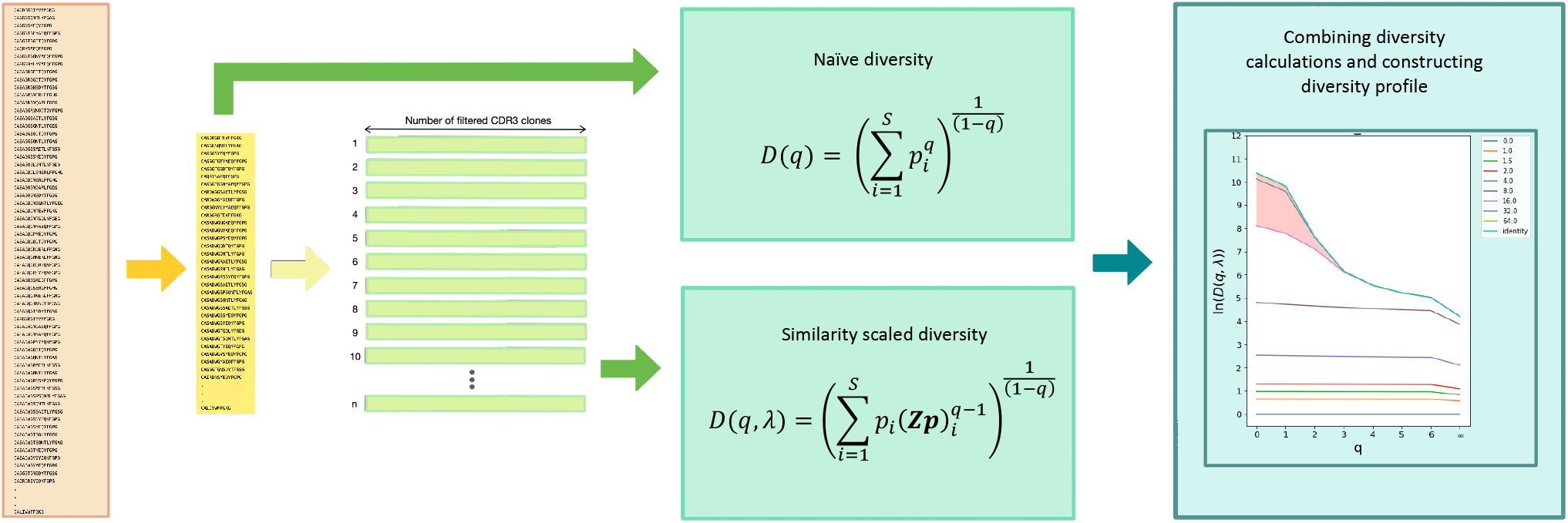
An overview of the TCRDivER algorithm. **I**.*Input*. The algorithm takes as input of CDR3 clone sequences with their clone sizes expressed as count or frequency. **II**.*Filtering*. The input sample is filtered to contain only “In” frame CDR3 regions. Afterwards, the repertoire is sampled for CDR3s randomly, based on the frequency (count) distribution in the original repertoire. The default subsampling size is 50000 CDR3 sequences, therefore the filtered and subsampled repertoires contain ≤ 50000 unique CDR3s. The CDR3 sequence counts in the subsampled repertoire are transformed into frequencies, so that the final output is a list of unique CDR3s with their respective frequencies summing up to 1 within the repertoire. **III**.*Calculating distance matrix* The filtered and downsampled repertoire is then provided as input for calculating the distance matrix. This step is split up so the original list of unique CDR3 sequences is divided into sublists of equal length which are in turn pairwise compared against the whole list of CDR3s using the BLOSUM45 alignment score (see Figure 6). The output is a set of files which contain portions of the distance matrix. If concatenated they would form the complete distance matrix. However, manipulating such a large file would computationally expensive. **IV**.*Calculating similarity scaled diversity* This step is split into two parts: calculating naive and similarity scaled diversity (see sections *Calculating Naive Diversity* and *Calculating Similarity scaled Diversity*. The first script takes in only the filtered and subsampled list of CDR3s and their respective frequencies, since *D*(*q*) is not dependent on CDR3 sequence distances. The output of this calculation is a .tsv file containing values of diversity at different values of *q*. The second script takes in both the list of CDR3s with their frequencies and the distance matrix “chunk” files. The calculation is done in a parallel fashion to reduce computational time. As output a .tsv file is given containing the values of calculated diversity at different values of *q* and *λ*. **V.** *Combining diversity calculations* The final step is joining the two diversity calculations into a complete overview of the diversity. Two previously obtained .tsv files with calculated diversity values are combined into one file containing all calculated values of diversity. Downstream this file is used for constructing the diversity profiles and further statistical analysis.

### Downstream analysis of TCRDivER output

The final output of TCRDivER is a table containing all the values of diversity calculated with a range of *q*s and *λ*s. The full downstream analysis is summarised in a Python3.6 jupyter notebook ***Kluyver et al. (2016)*** available alongside the main TCRDivER algorithm at https://github.com/sciencisto/TCRDivER. The diversity profiles were constructed using the seaborn package ***Waskom et al. (2017)***. Areas between *λ* curves of ln (*D*(*q, λ*)) was calculated within the numpy framework ***Oliphant (2006)*** as an integration using the composite trapezoidal rule. Average Δln(*D*(*q, λ*)) for small *λ*s was calculated for *λ* = {0.0, 0.1, 0.2, 0.3, 0.4, 0.5} as the mean of the average difference between ln (*D*(*q, λ*)) at each calculated *q*. Slopes of diversity when *q* = 0 → 1, *q* =1 → 2 and *q* = 0 → 2 were calculated as differences between the diversities when *q* = 0, 1 and 2. We are aware that this is not the slope of diversity at the specific values of *q*, as we also provide the mathematical evaluation of the slope at these timepoints (See Appendix 1 Evaluating the slope at *q* = 1). However, due to time and memory considerations we have opted for the simplified calculation as we feel that it represents the slope sufficiently. For future implementations, we will update the algorithm to include the analytical slope evaluation. Principal components analysis was performed as implemented in the scikit-learn package ***Pedregosa FABIANPEDREGOSA et al. (2011)***.

## Acknowledgements

We would like to thank Prof. Nir Friedman, Dr. Shlomit Reich-Zeliger and Dr. Erik Shifrut from the Weizmann Institute, Rehovot, Israel for generating and sharing the TCR sequence data analysed in this paper.

## Appendix 1 Evaluation of naive diversity of the first order

We start with:

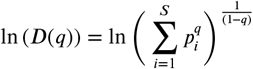

Exploring the limit as *q* approaches 1 allows us to apply L’Hopitals rule:

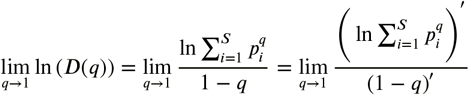

The solution is:

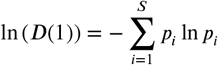

This is equivalent to:

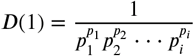

### Evaluating the slope at *q* = 1

We start with the definition of^*q*^ *D*:

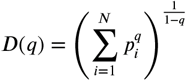

define *S* as the internal sum

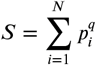

Now we can write

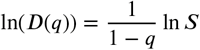

Let’ denote differentiation with respect to *q*. Now evaluate

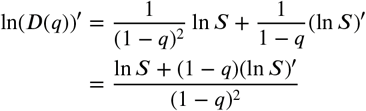

To evaluate the limit as *q* goes to 1 we need to apply l’hopital’s rule twice. Calling the numerator *t*

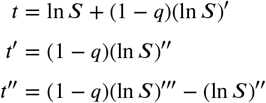

Since ln *S* and all its derivatives are finite as *q* goes to 1

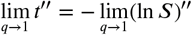

Call the denominator *b*

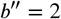

and

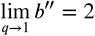

Putting this together we have

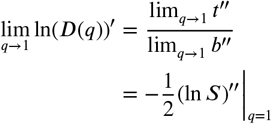

We need

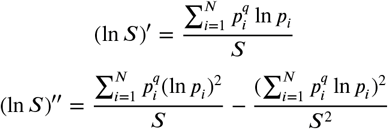

Because *p_i_* is a probability distribution

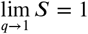

which means (since the limit of quotient is the quotient of the limits)

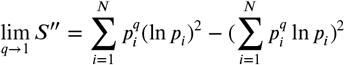

Finally, we want to evaluate *D*(*q*)′ and then take the limit as *q* goes to 1.

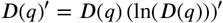

and

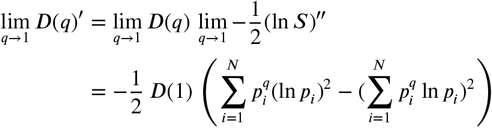

To generalise to the case **Z** ≠ **I** we simply have to replace *S* with

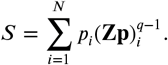

In this case

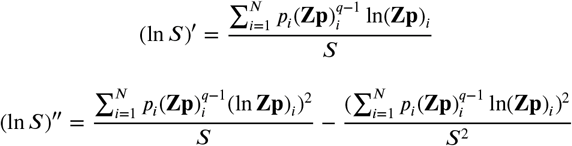

### Evaluation of similarity scaled diversity of the first order

We start with:

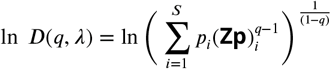

Rewritting the equation, calculating the limit as q approaches 1 and applying L’Hopitals rule:

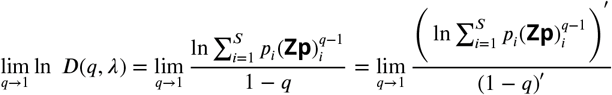

The result is:

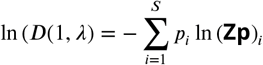

which is equivalent to:

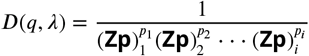

### Evaluation of naive diversity of the infinity order

We start with the formula for naive diversity and extract the largest clone frequency *p_max_*:

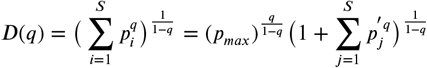

where 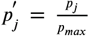 for *j* ≠ *max*, and *p_max_* is represented in the first term of the sum. Since a limit of products is a product of limits, it follows:

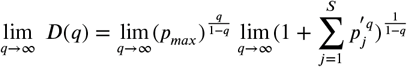

The first limit is evaluated as:

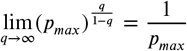

The second limit is evaluated by taking the logarithm:

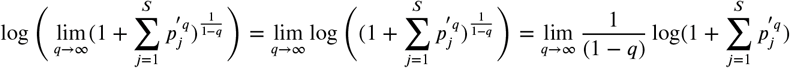

Since 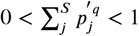, the bounds of logarithm are:

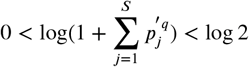

which gives:

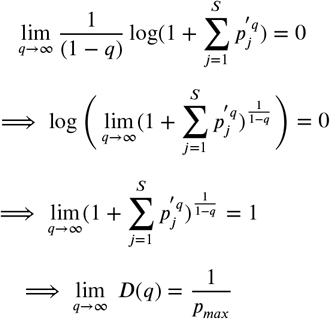

### Evaluation of similarity scaled diversity of the infinity order

We start with:

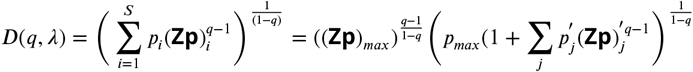

where the term that has been pulled out is the one for which (**Zp**)_*i*_, is maximum. The *p_max_* is the corresponding *p_i_*. As before, the 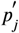 are defined as 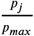 for *j* ≠ *max* and 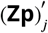 is defined as 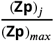. Again, the limit splits in to two factors:

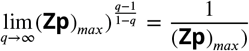

Taking the log of the second term gives:

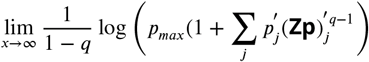

and now the log is bounded by:

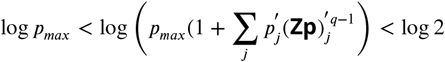

so again the limit of the log second factor in (*) is 0, and limit of the factor itself is 1. The end result is:

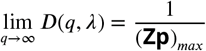

which reduces to the correct limit when **Z**=**I** which is the naive diversity.

## Appendix 2 Evaluation Δ*ln*(*D*(*q, λ*)) for small *λ*: Perturbation around *λ* = 0

Conjecture: gradient of 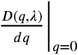 is a decreasing function of *λ*. N.B. *D*(*q, λ*) is an increasing function of *λ* for all *q*.

We start with the assumption that for *λ* around 0:

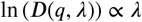

Where ln(*D*(*q, λ*)) is:

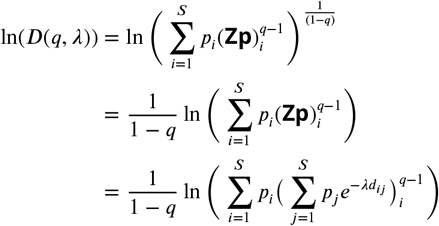

For *λ* → 0 by applying Taylor expansion 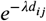 reduces to 1 – *λd_ij_* which gives:

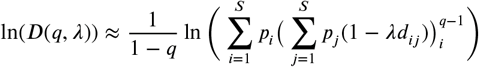

We can then rewrite:

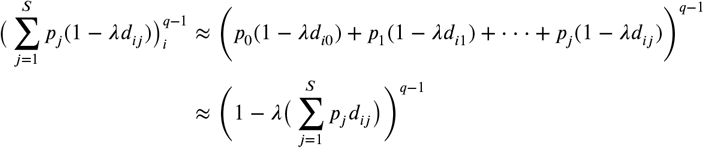

By applying the binomial expansion we arrive at:

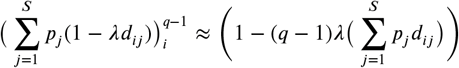

By substituting the derived expressions in the formula for ln (*D*(*q, λ*)) and keeping in mind that 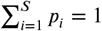, we can write:

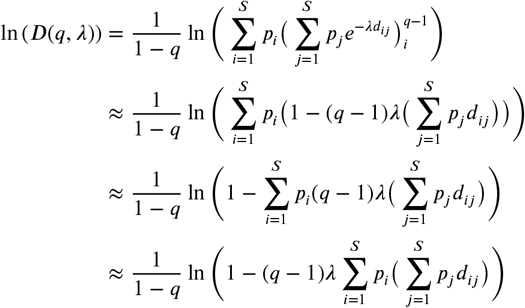

By applying the linear approximation *ln*(1 – *x*) ≈ *x*, we finally arrive:

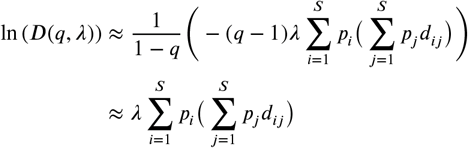

Note that the final form of the evaluation of *D*(*q, λ*) for *λ* → 0 is independent of the order of diversity *q*. It is solely dependent on the distance between CDR3 sequences weighted by their respective frequencies.

### Evaluation Δ*ln*(*D*(*q, λ*)) for small *λ* and it’s relationship to distance

By evaluating Δ*ln*(*D*(*q, λ*)) for two values of small *λ*, where *λ*′ > *λ*″ we arrive at:

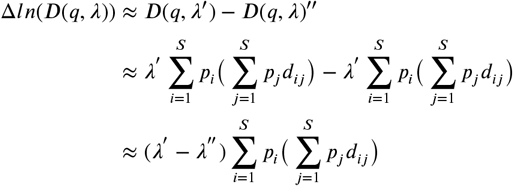

It is evident that Δ*ln*(*D*(*q, λ*)) is linearly dependent on the distances between CDR3s and their probabilities. In the case of two hypothetical repertoires, **I** and **II**, which have a uniform distribution of CDR3 frequencies within the repertoire 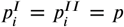 and distances between CDR3s 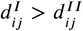, Δ*ln*(*D*(*q, λ*)) for repertoire **I** is larger than Δ*ln*(*D*(*q, λ*)) for repertoire **II**. That is with the increase of similarity between CDR3s, the area between the curves for small *λ*s decreases (Appendix 2 Figure 1 C.). Alternatively, if the distances between CDR3s of the two repertoires are the same 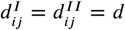, and the distribution is still uniform, but the number of clones differs so that repertoire **I** has less clones than **II** i.e. 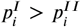, then Δ*ln*(*D*(*q, λ*)) is larger than Δ*ln*(*D*(*q, λ*)). Meaning that repertoires with more abundant clones have a larger Δ*ln*(*D*(*q, λ*)) for small *λ*s.

### Evaluation Δ*ln*(*D*(*q, λ*)) for larger *λ*s and it’s relationship to distance

In order to evaluate the relationship between CDR3 clone distance and the area between the curves of larger *λ*s we have constructed three mock repertoires. The reperotires constist of 100 CDR3s that are uniformly distributed in the repertoire, i.e. 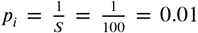. For each mock repertoire a mock distance matrix was calculated so that the distance between the CDR3s within the repertoire were equal, but that they differ between the repertoires. The distances were 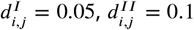 and 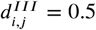, for repertoire I, II and III respectively when *i* ≠ *j*, else *d_i,j_* = 0 for *i* = *j*. Individual *λ* curves of the diversity profiles straight lines - a remnant of uniform distribution of CDR3 frequencies in the repertoire (Appendix 2 Figure 1).

**Appendix 2 Figure 1.**
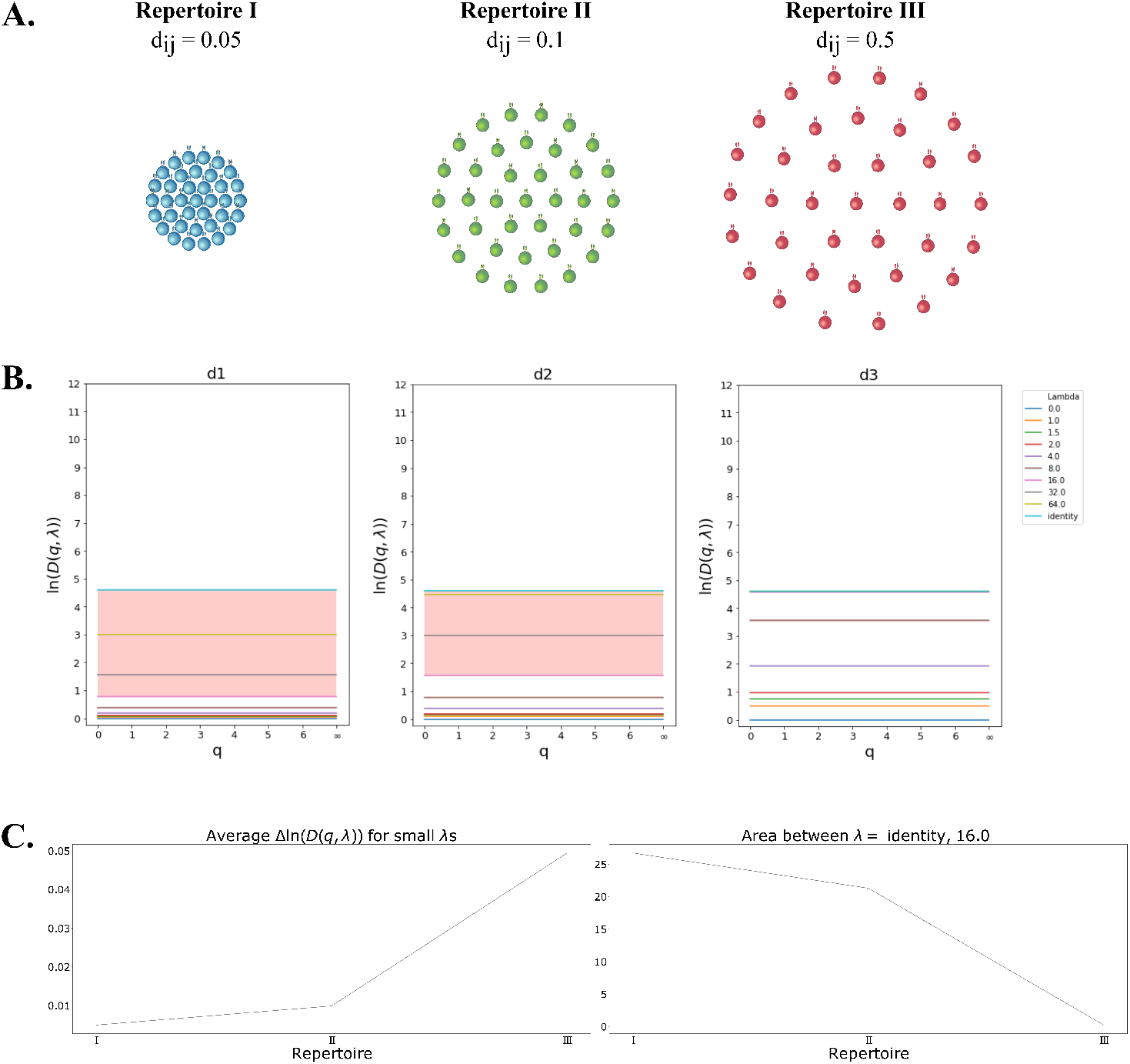
Effect of CDR3 distance shown in three mock repertoires with a uniform distribution of 100 CDR3 clones in in the repertoire. **A.** Schematic representation of the three mock repertoires with the distances *d_ij_* between CDR3s increasing from repertoire I to III. **B.** Diversity profiles calculated based on the probability distribution and *d_ij_* for CDR3s in the mock repertoires. The frequency of seeing each CDR3 clone in all the repertoires, since they consits of 100 uniformly distributed CDR3s, is 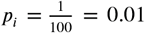 **C.** Calculated values of average Δ ln (*D*(*q, λ*)) for small *λ*s and calculated area between *λ* identity and 16 curves for the three repertoires, shown left to right respectively.

## Supplementary information

### 1 Murine Dataset

Results concerning the murine CD4^+^ TCR repertoire dataset following immunisation with Complete Freund’s Adjuvant (CFA) with or without the additon of Ovalbumin (OVA) antigen. The dataset also contains unimminused mice. The samples were collected at 3 timepoints (5, 14 and 60 day) for immunised mice, and day 0 for unimmunised mice. Two TCR distance metrics were used a BLOSUM45 and Atchley factor based score. Each subsection will therefore be marked with the distance metric

#### 1.1 Diversity Profiles-BLOSUM45

Diversity profiles calculated for the murine dataset with 50000 subsample size. The profiles are organised into tables according to post-immunisation sample collection time with untreated mice, day 5, day 14 and day 60 in Table 1, 2, 3 and 4, respectively.

**Table 1:**
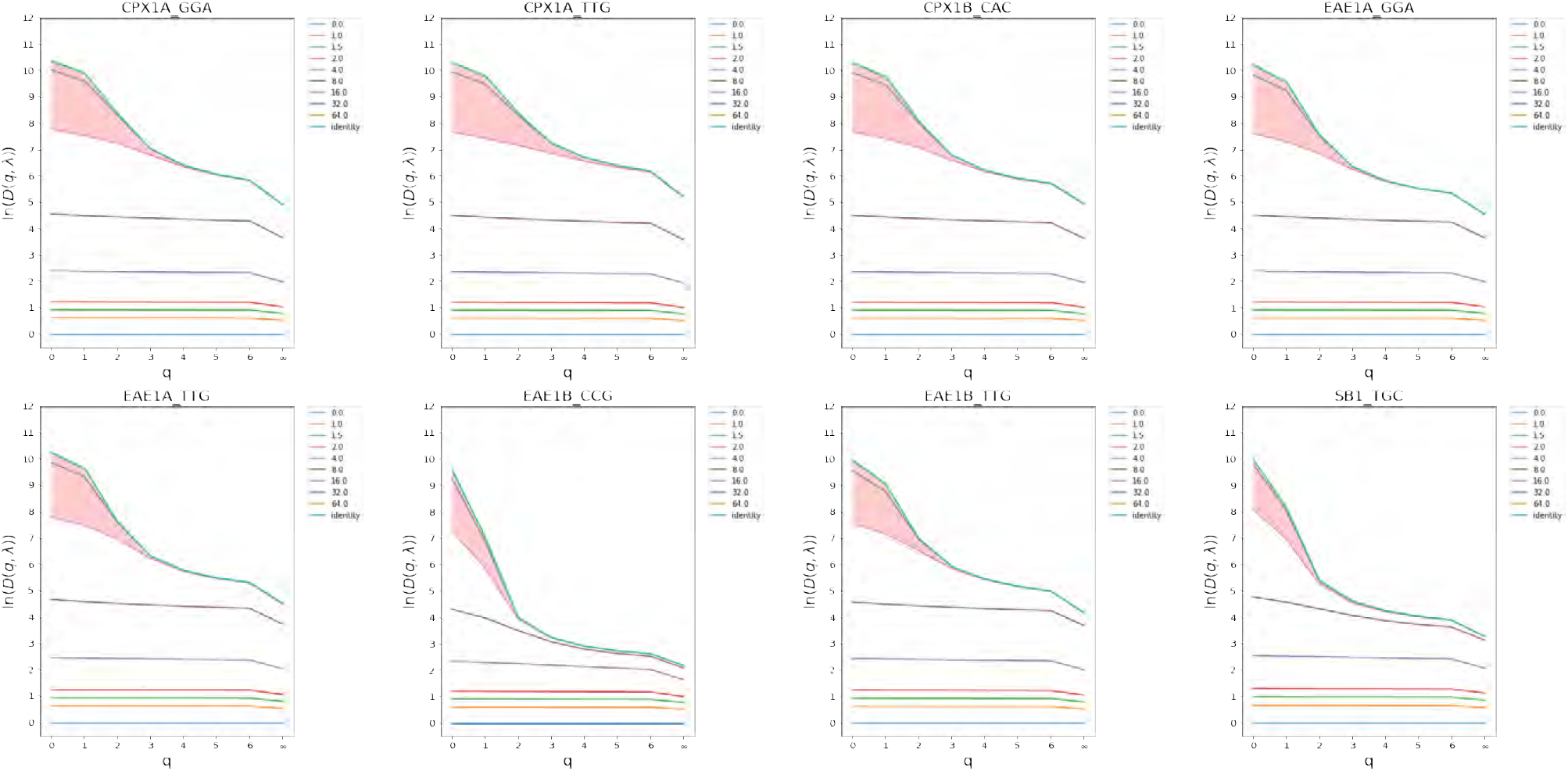
Untreated day 0

**Table 2:**
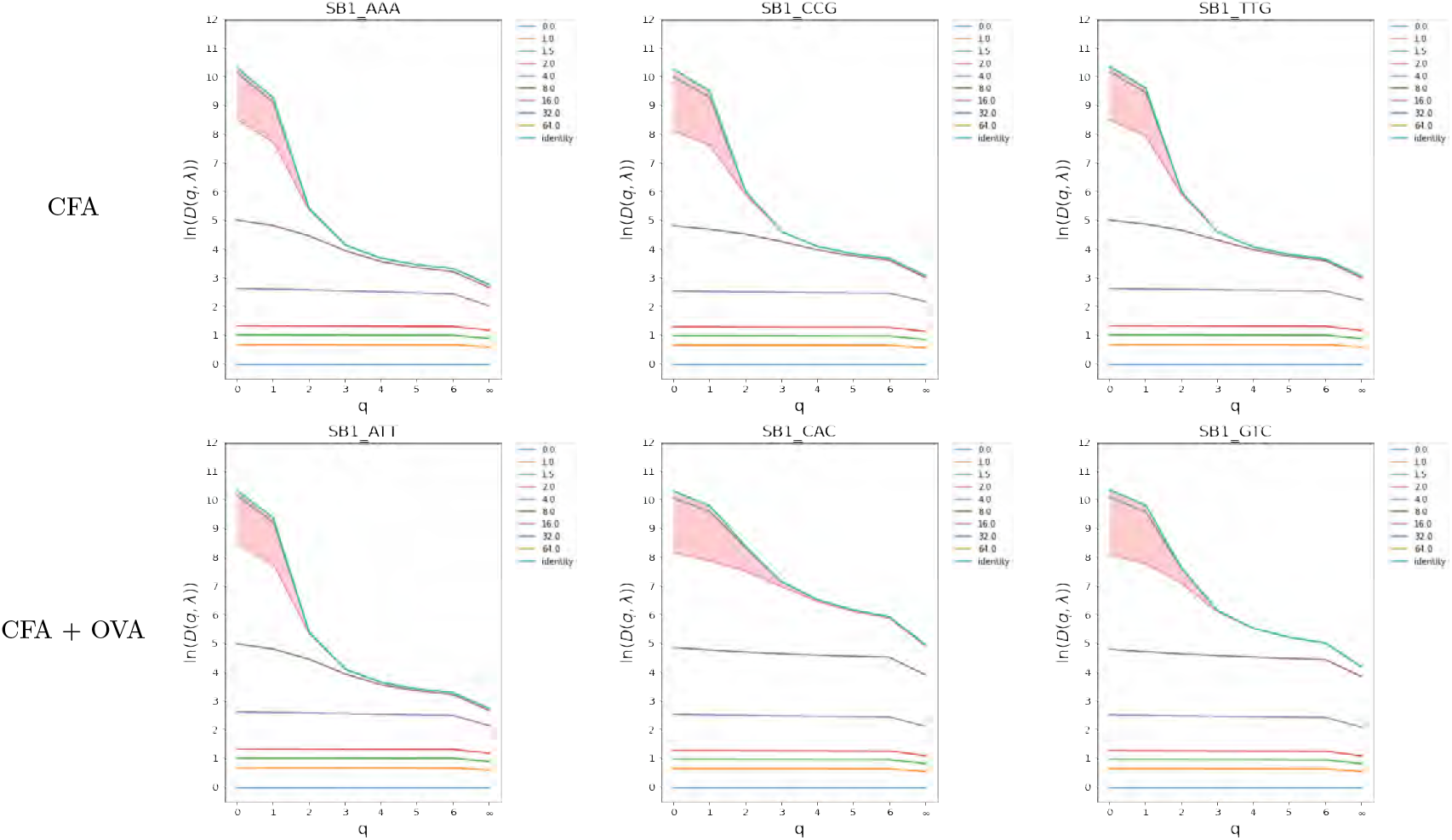
Immunised day 5

**Table 3:**
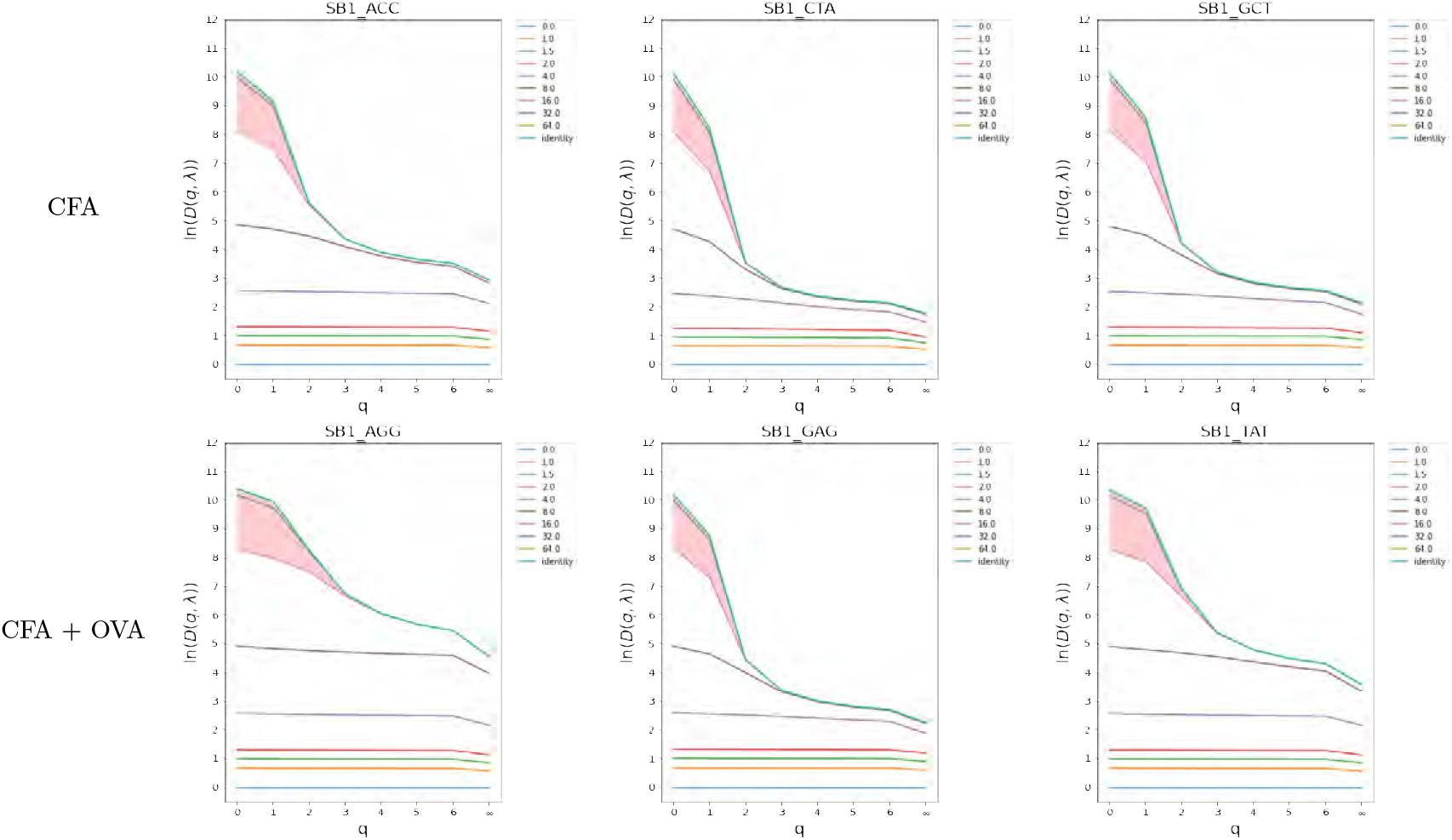
Immunised day 14

**Table 4:**
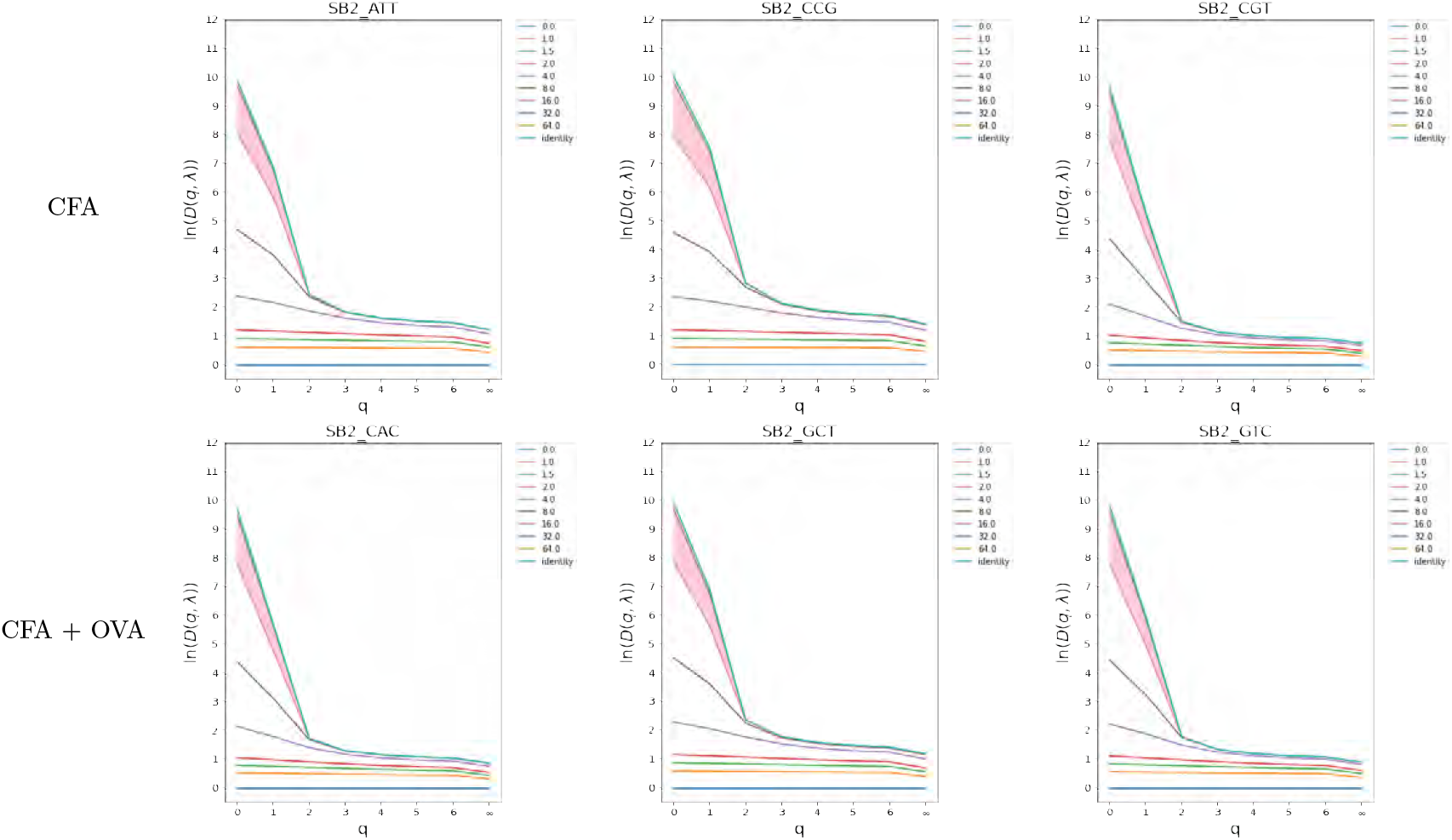
Immunised day 60

#### 1.2 Naive (*q* = 0^9^)^9^ diversity profiles-BLOSUM45

**Figure 1:**
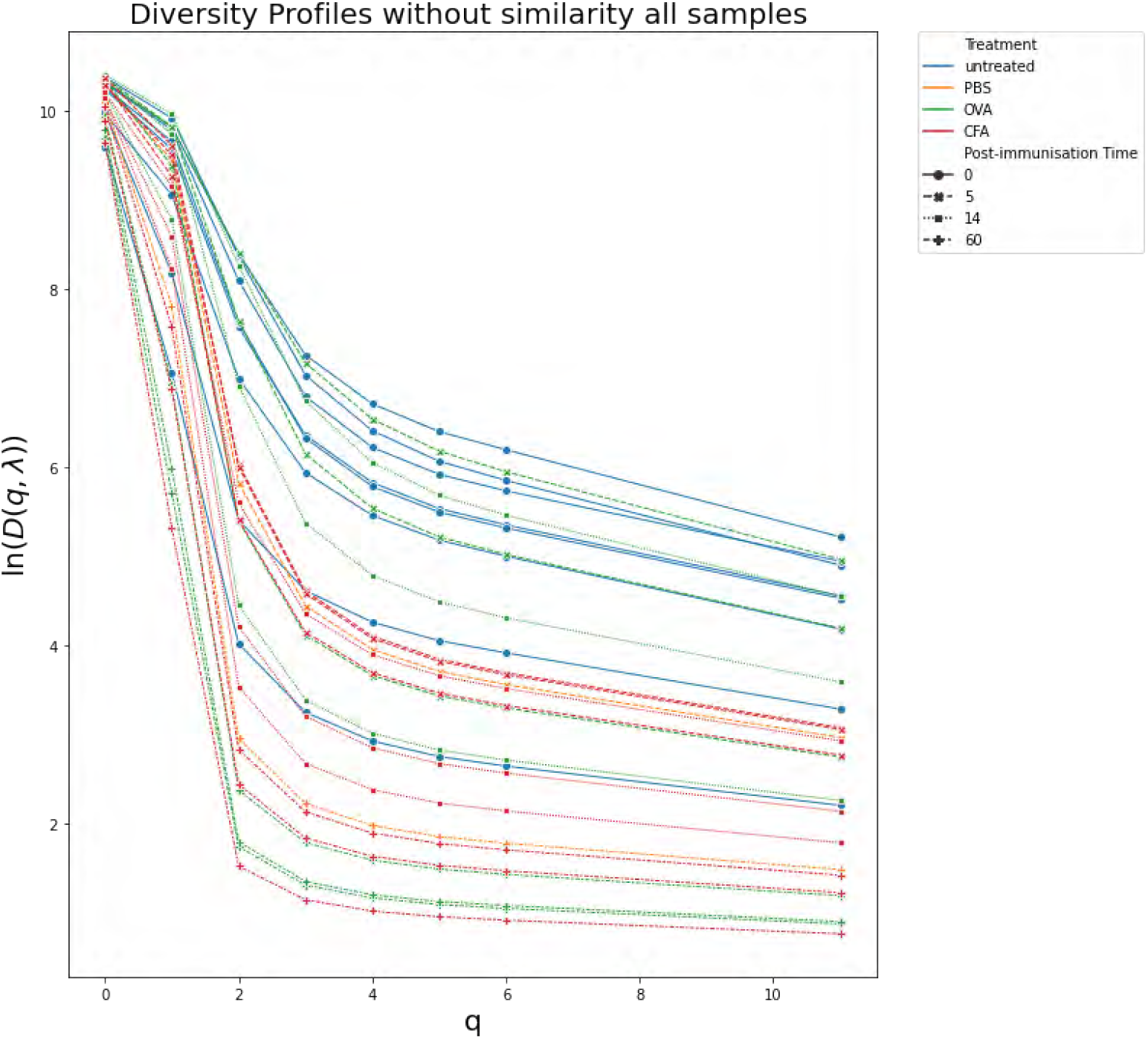
Naive (*q* = 0) diversity profiles plotted for all murine samples. Frequent crossings of the curves can be observed illustrating that the rank order of samples depends on the specific choice of index.

#### 1.3 PCA on natural logarithm transformed values of true diversity-BLOSUM45

**Figure 2:**
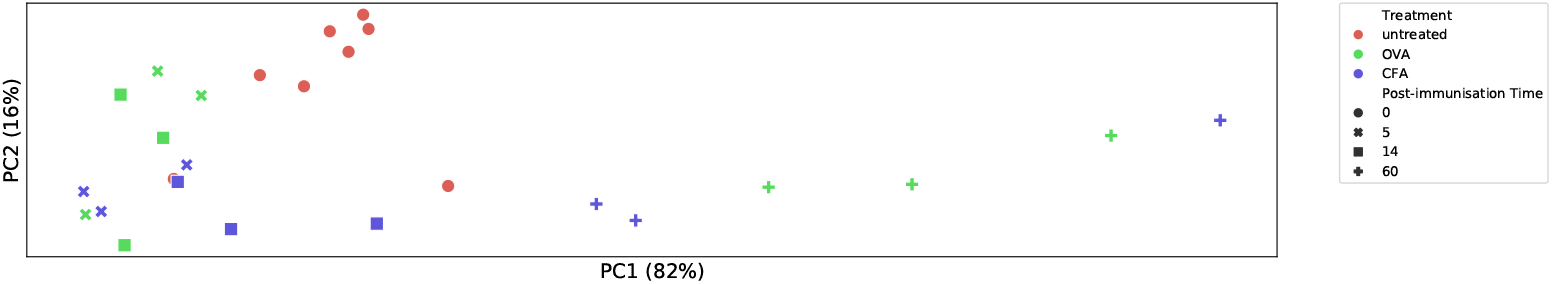
PCA on values of true diversity *D*(*q*, λ). for the murine dataset. The aspect ratio corresponds to variation found by PCA.

#### 1.4 PCA on diversity values from the randomised murine dataset with random frequencies-BLOSUM45

**Figure 3:**
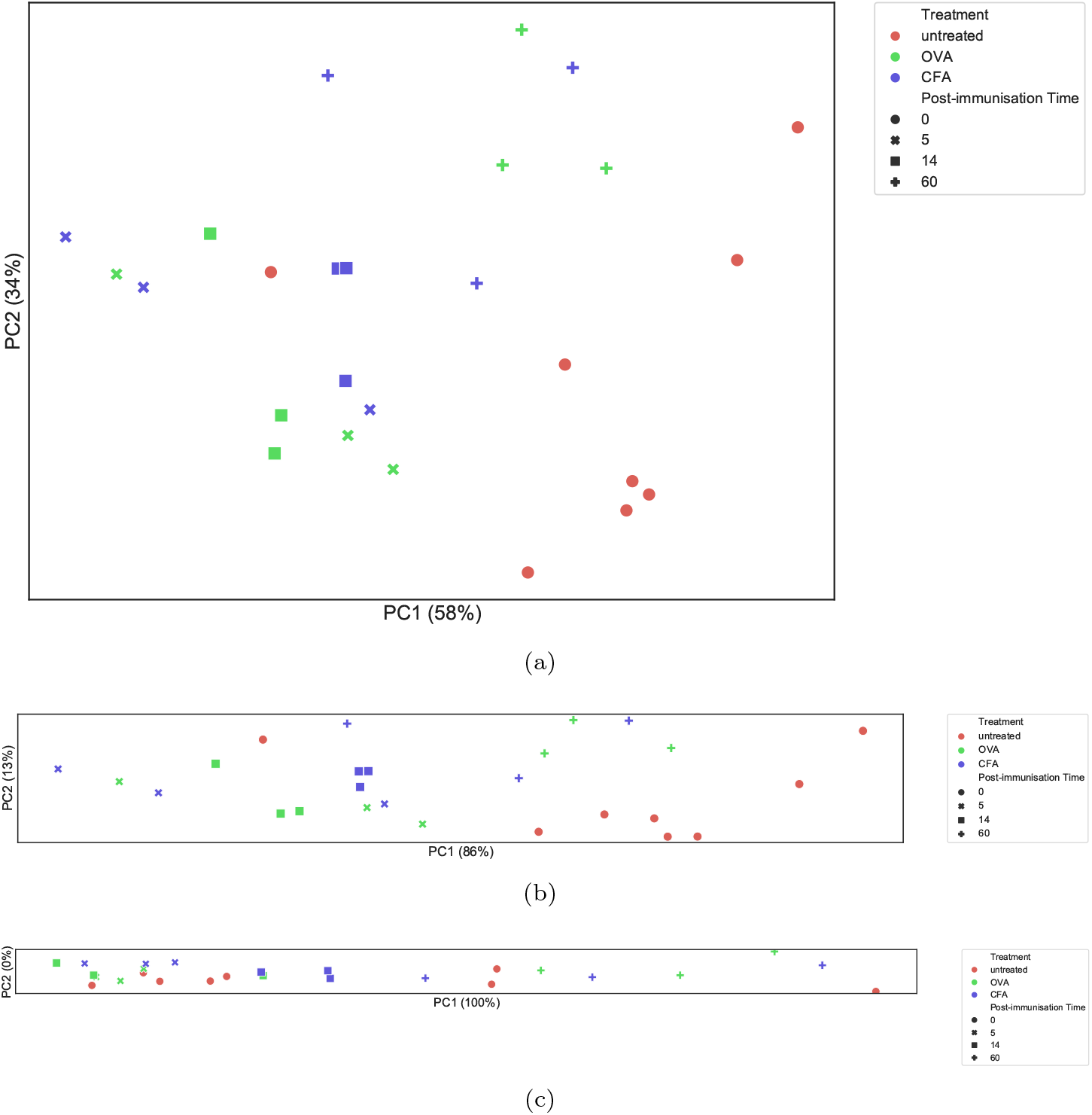
Principal Components Analysis on diversity calculated for the randomised murine dataset. The aspect ratio corresponds to variation found by PCA. **a.** PCA on features extracted from the diversity profiles constructed from the true diversity *D*(*q*, λ). **b.** PCA on values of true diversity *D*(*q*, λ). **c.** PCA on naive diversity values *D*(*q*), i.e. λ = identity.

#### 1.5 Diversity Profiles-Atchley factor distance

Diversity profiles calculated for the murine dataset with 50000 subsample size. The profiles are organised into tables according to post-immunisation sample collection time with untreated mice, day 5, day 14 and day 60 in Table 1, 2, 3 and 4, respectively.

**Table 5:**
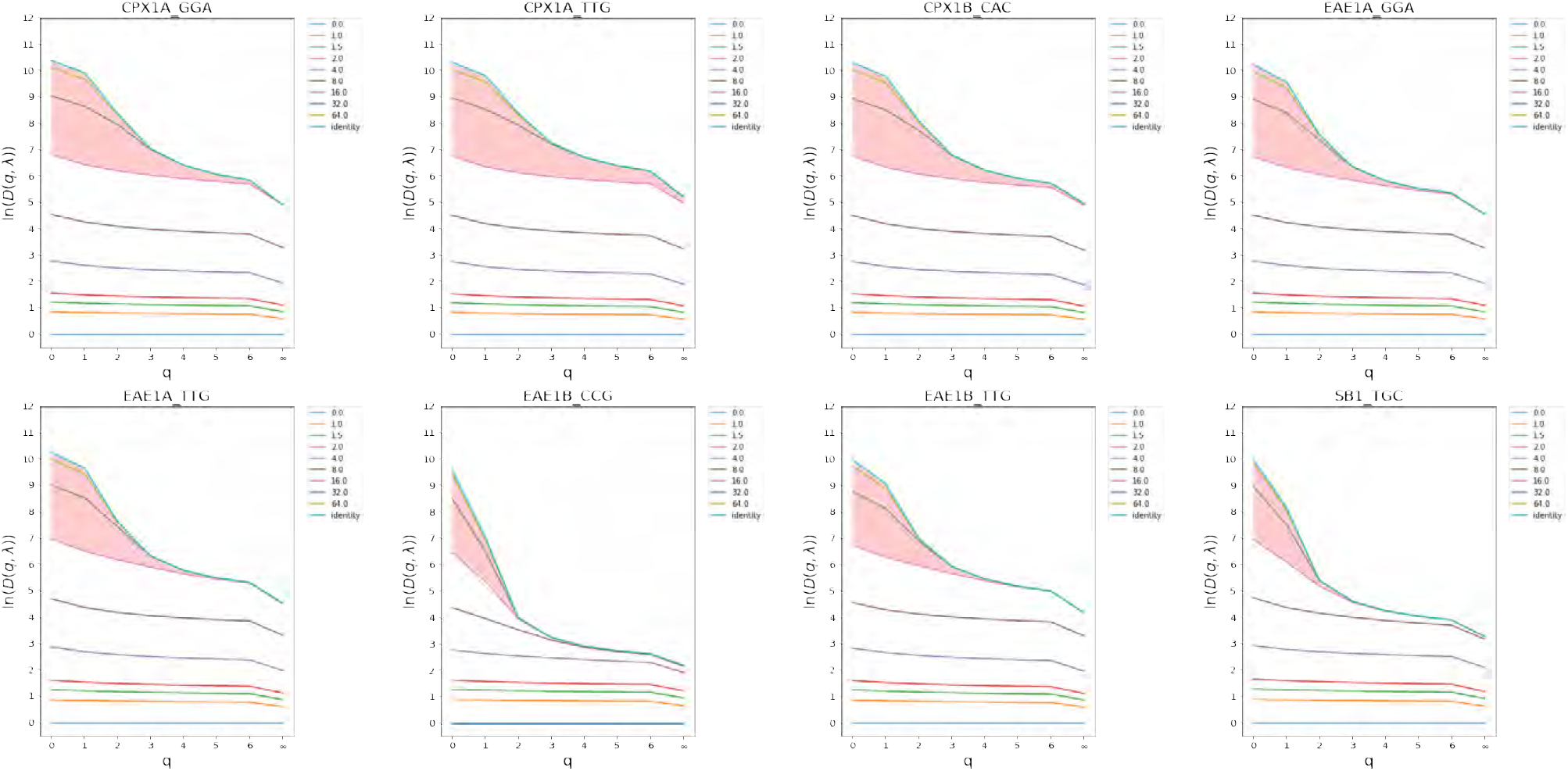
Untreated day 0

**Table 6.**
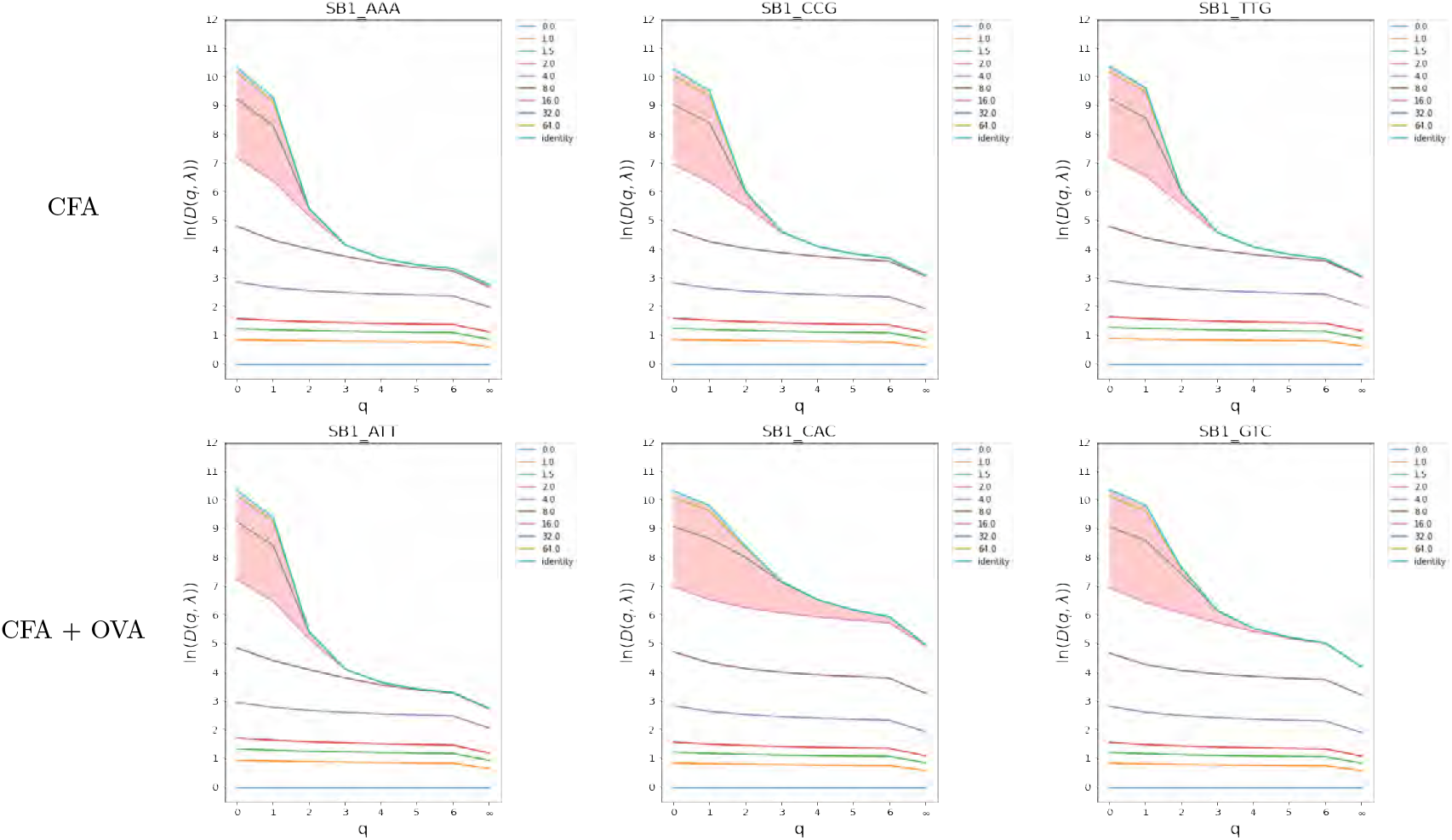
Immunised day 5

**Table 7:**
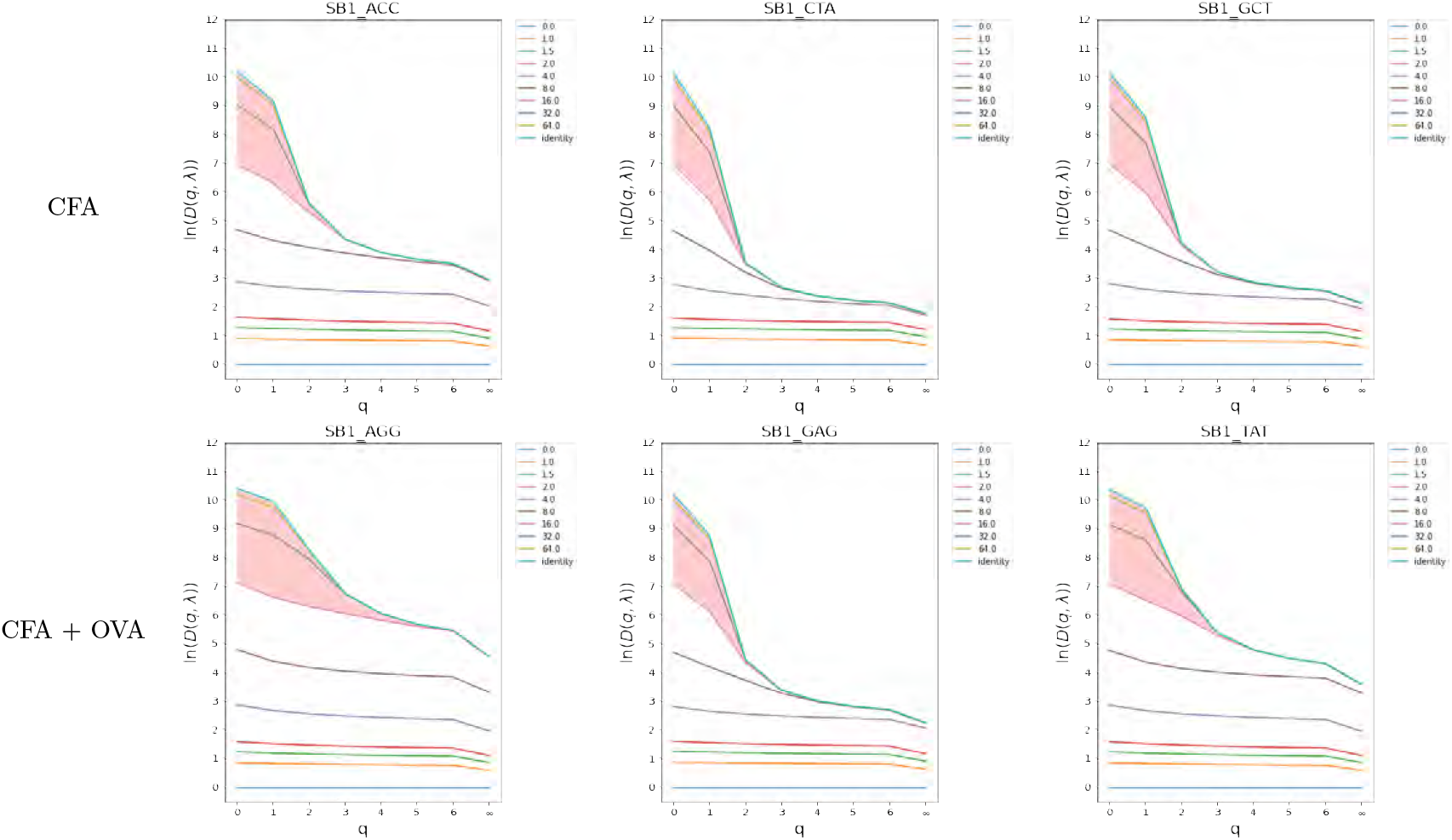
Immunised day 14

**Table 8:**
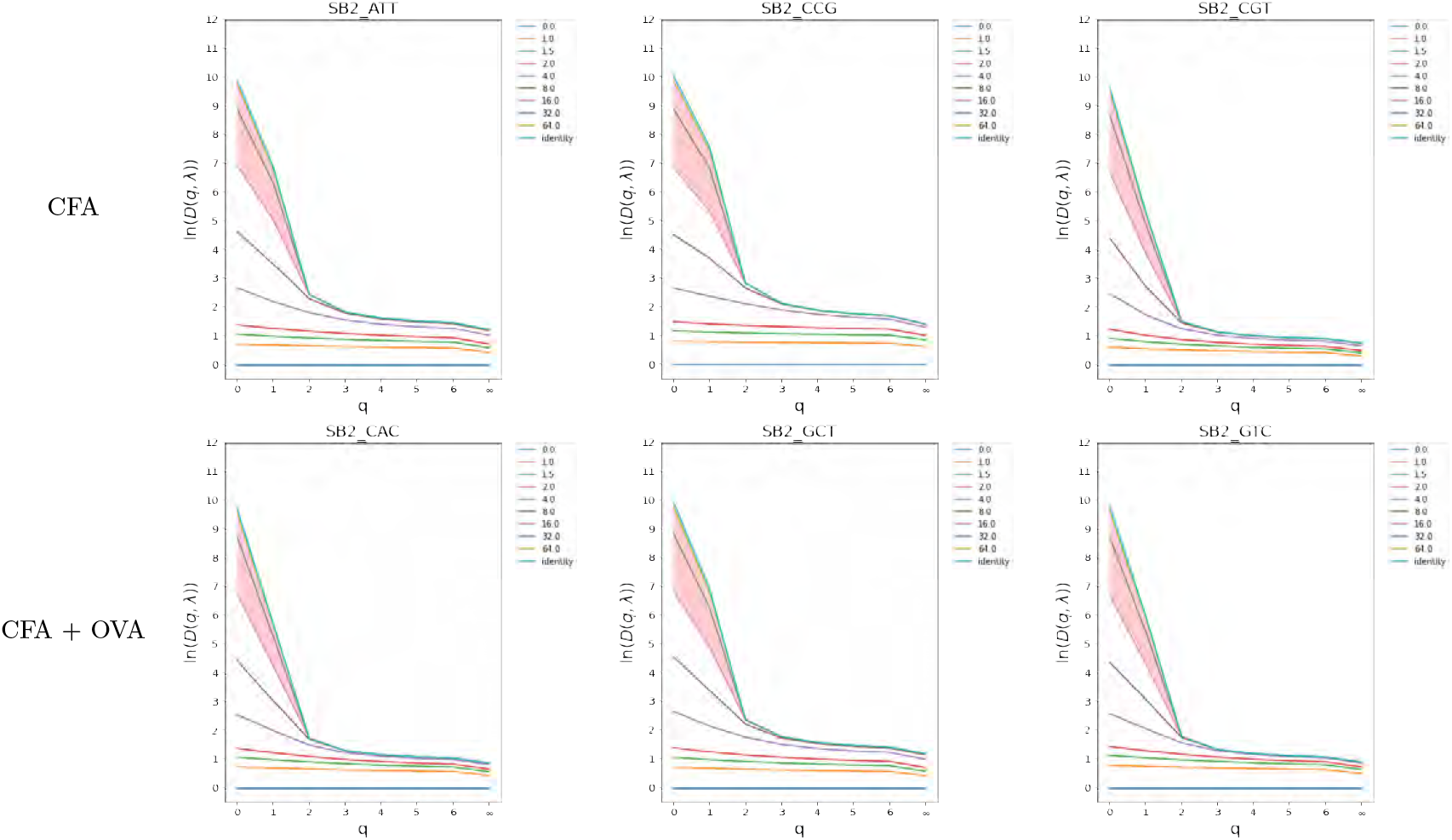
Immunised day 60

#### 1.6 Naive (*q* = 0^9^)^26^ diversity profiles–Atchley factor distance

**Figure 4:**
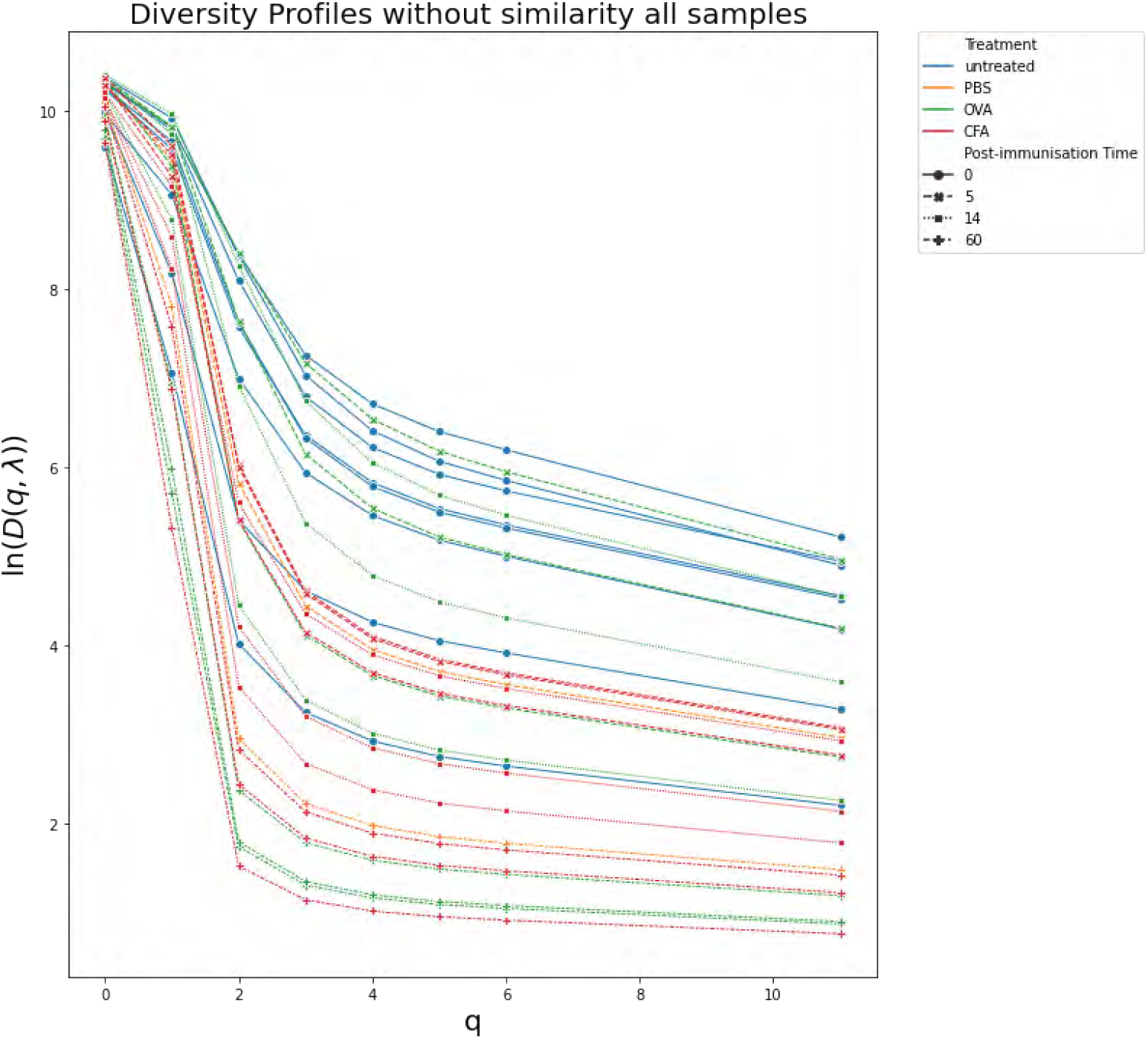
Naive (*q* = 0) diversity profiles plotted for all murine samples. Frequent crossings of the curves can be observed illustrating that the rank order of samples depends on the specific choice of index.

#### 1.7 PCA on natural logarithm transformed values of diversity–Atchley factor distance

**Figure 6:**
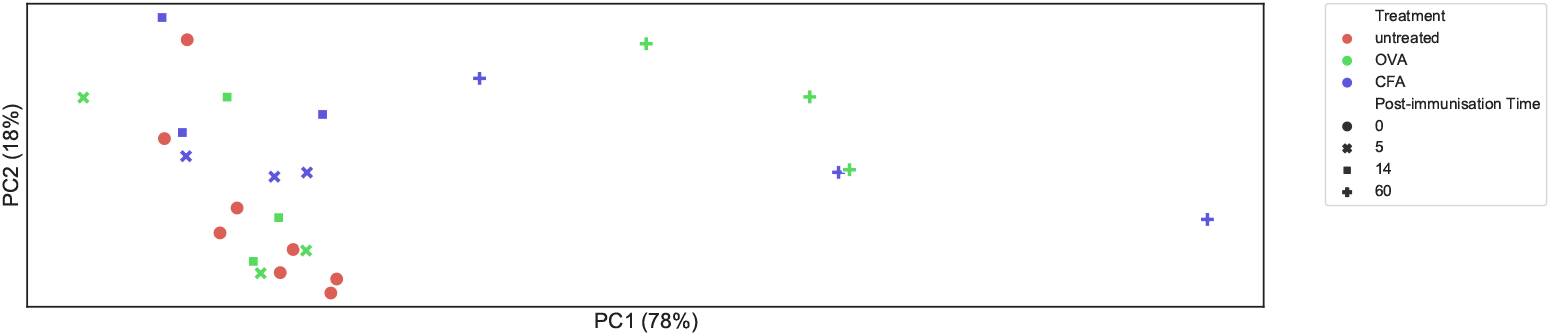
PCA on values of true diversity *D*(*q*, λ). for the murine dataset. The aspect ratio corresponds to variation found by PCA.

**Figure 5:**
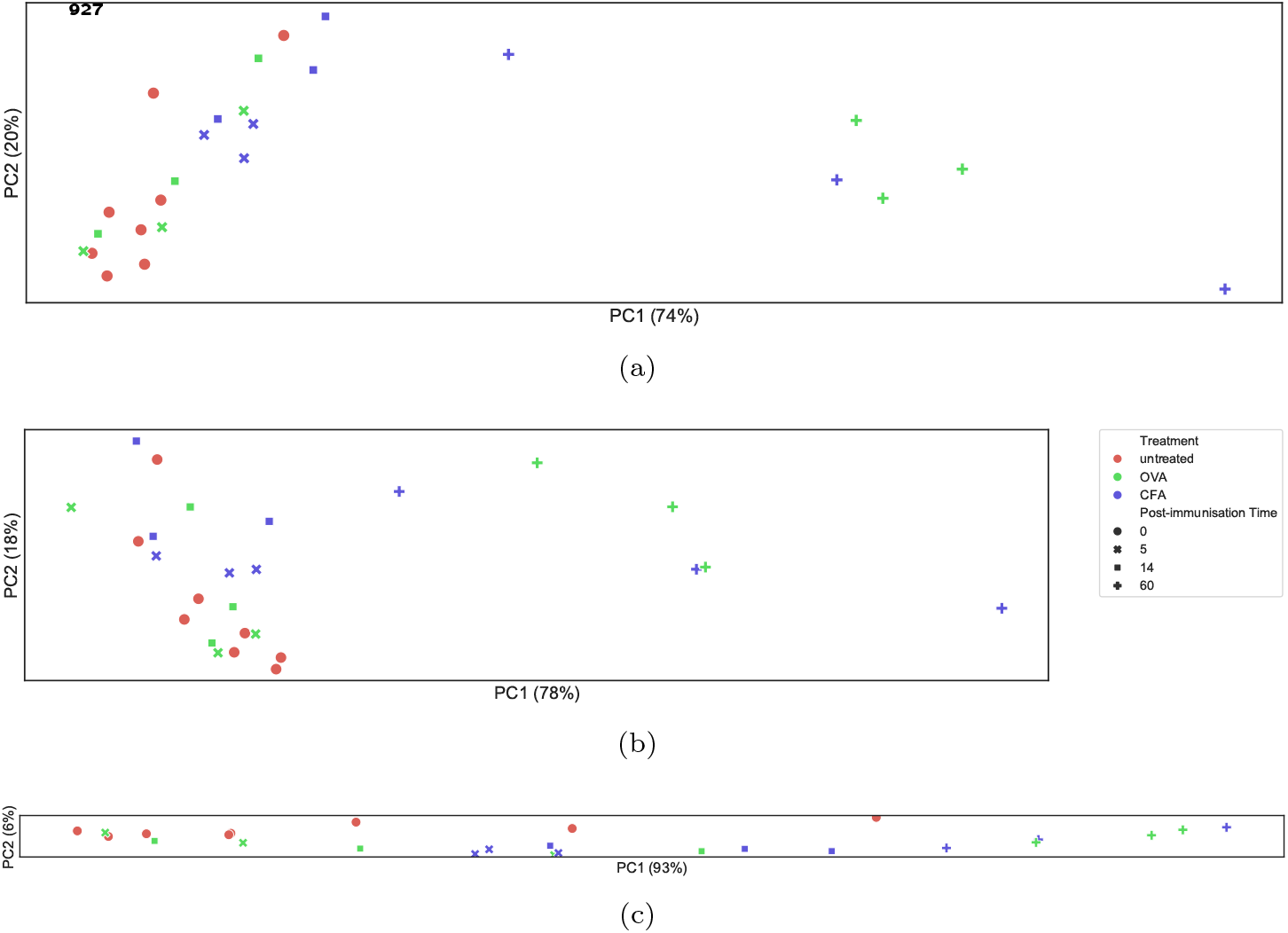
Principal Components Analysis on diversity calculated for the murine dataset using the Atchley Factor distance for TCRs. The aspect ratio corresponds to variation found by PCA. **a.** PCA on features extracted from the diversity profiles constructed from the true diversity *D*(*q*, λ). **b.** PCA on values of true diversity *D*(*q*, λ). **c.** PCA on naive diversity values *D*(*q*), i.e. λ = identity.

#### 1.8 Trends of three features extracted from divPs versus timepoint and treatment regime-Atchley factor distance

**Figure 7:**
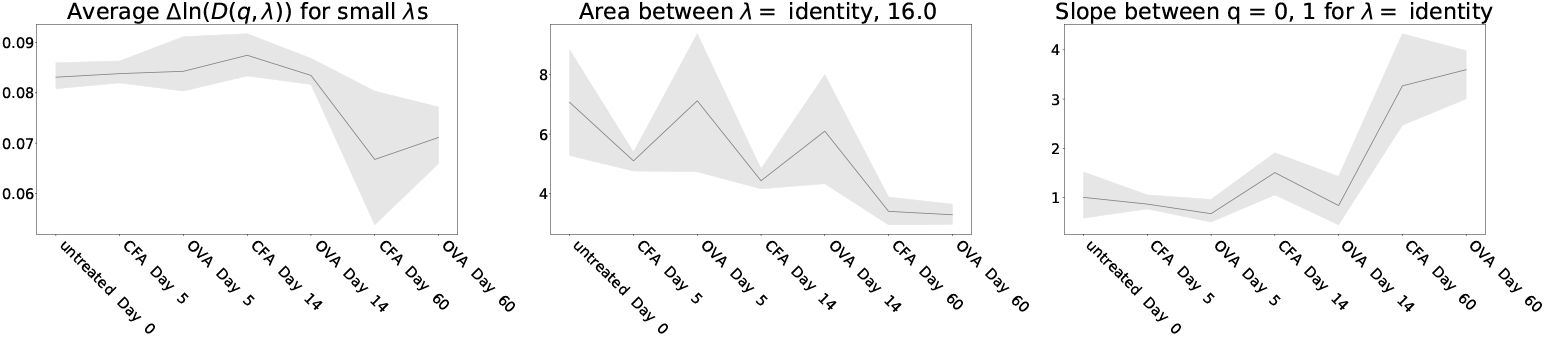
Trends of three features extracted from divPs are shown versus the treatment regime and timepoints ending with the latest timepoint. The features are, from left to right: average Δln*D*(*q*, λ) for small λs, between curves of λ = identity and 16.0 and slope of *q* = 0 → 1 for value of λ identity. The line connects the mean values of the features for all samples within a group and the shaded area represents the confidence interval.

#### 1.9 DivP features relationships-Atchley factor distance

**Figure 8:**
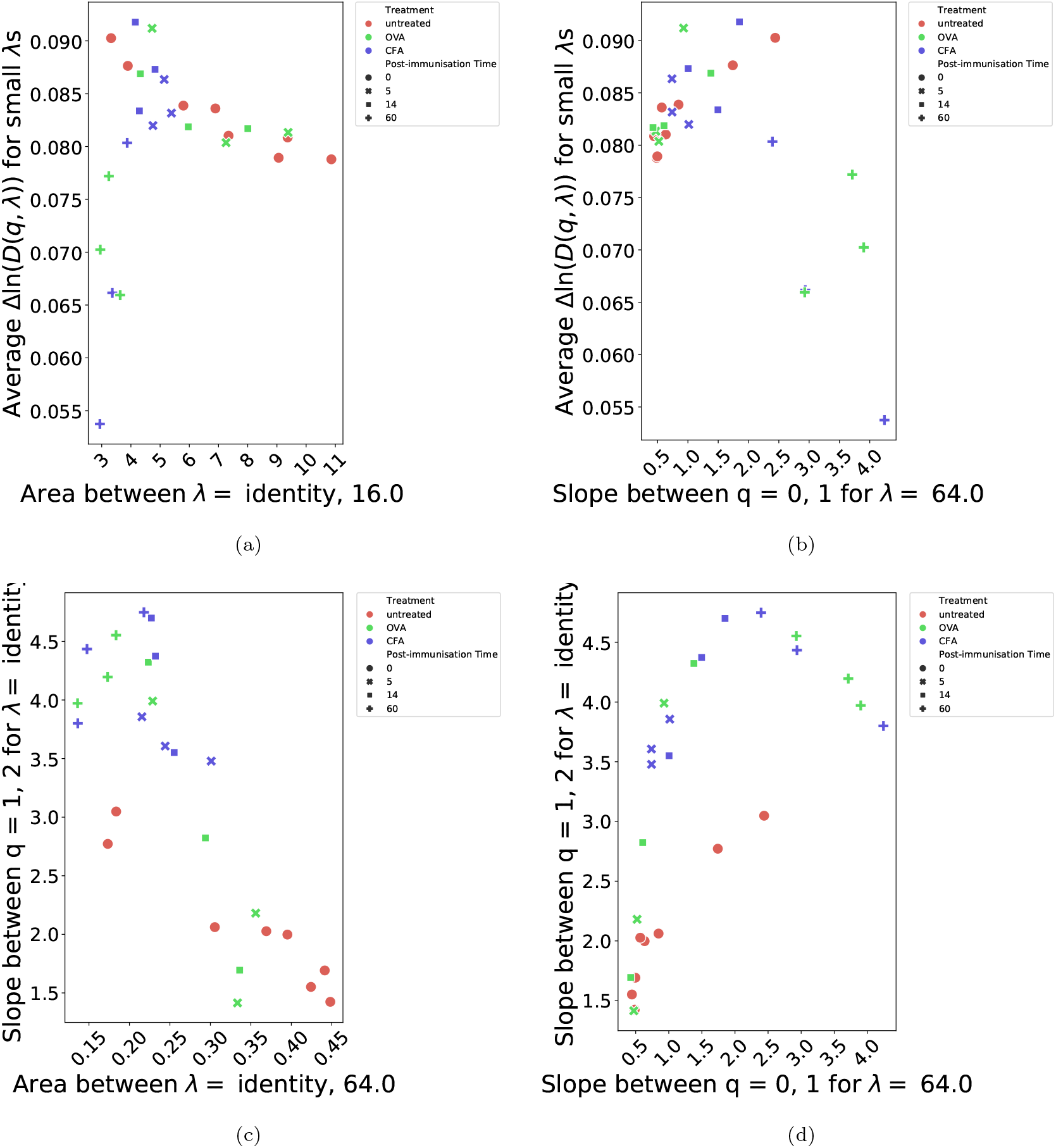
Graphs showing relationships between some of the divP features. **a.** average Δln *D*(*q*, λ) for small λs is shown versus the area between curves of λ = identity and 16.0; **b.** average Δln *D*(*q*, λ) for small λs is shown versus the slope of *q* = 0 → 1 for value of λ 64.0; **c.** slope of *q* =1 → 2 for value of λ identity (i.e. naive diversity) is shown versus the area between curves of λ = identity and d. 64.0; slope of *q* = 1 → 2 for value of λ identity (i.e. naive diversity) is shown versus the slope of *q* = 0 → 1 for value of λ 64.0.

### 2 Human Dataset

Results concerning the human TCR repertoire dataset following combination therapy (RT and anti-CTLA4 blockade ipilimumab. The patients were stratified into RECIST response criteria and by time of sample collection. We show the results in the same order.

#### 2.1 Diversity Profiles

Diversity profiles calculated for the human dataset with 50000 subsample size. The profiles are organised according to RECIST criteria and timepoint in Tables 5 to 12 for progressive disease (PD) day 0 and 22, stable disease (SD) day 0 and 22, partial responders (PR) day 0 and 22 and complete responders (CR) day 0 and 22, respectively. The distance metric used in estimating diversity was based on the BLOSUM45 alignment.

**Table 9:**
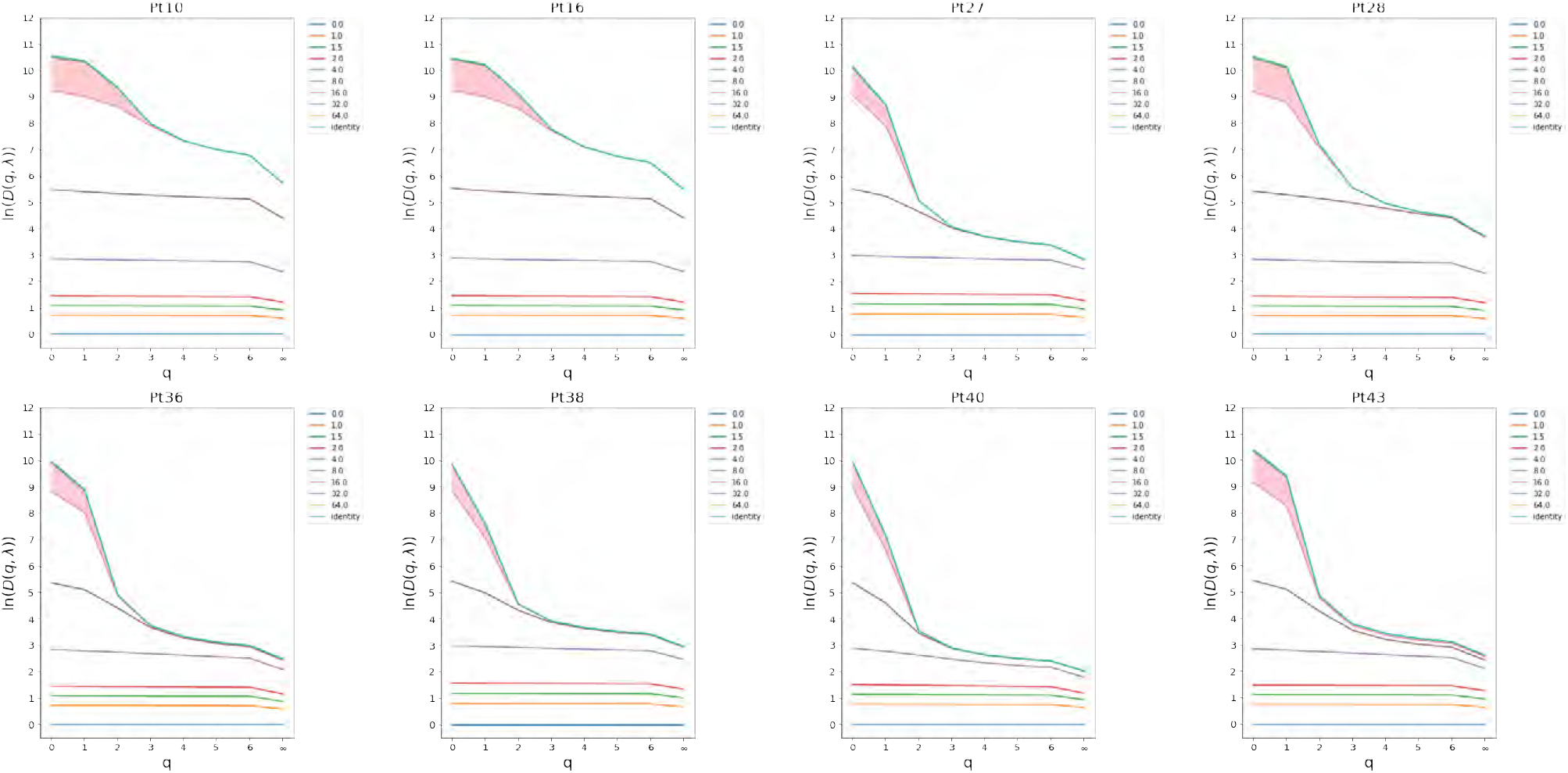
Progressive Disease (PD) Day 0

**Table 10:**
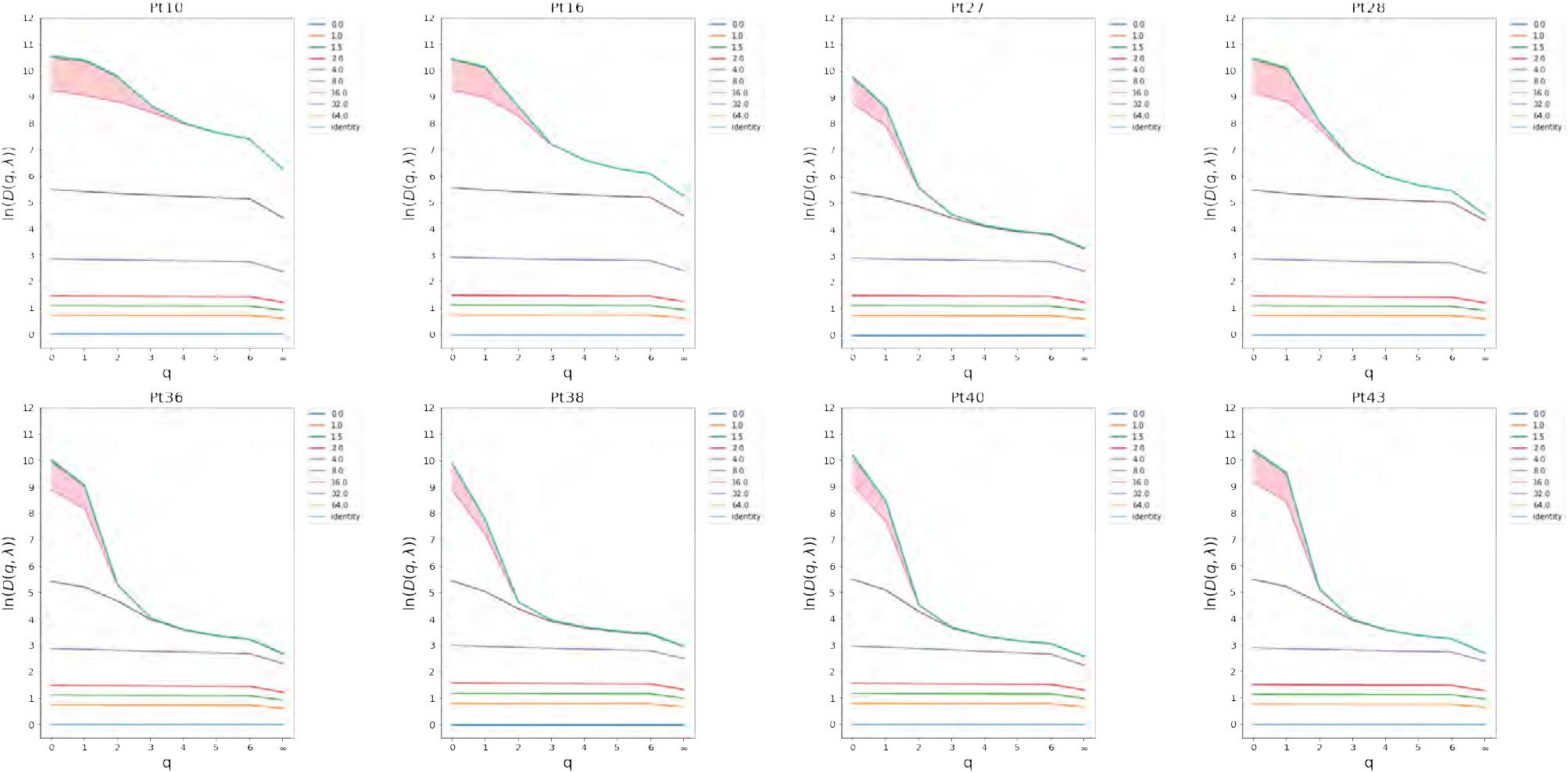
Progressive Disease (PD) Day 22

**Table 11:**
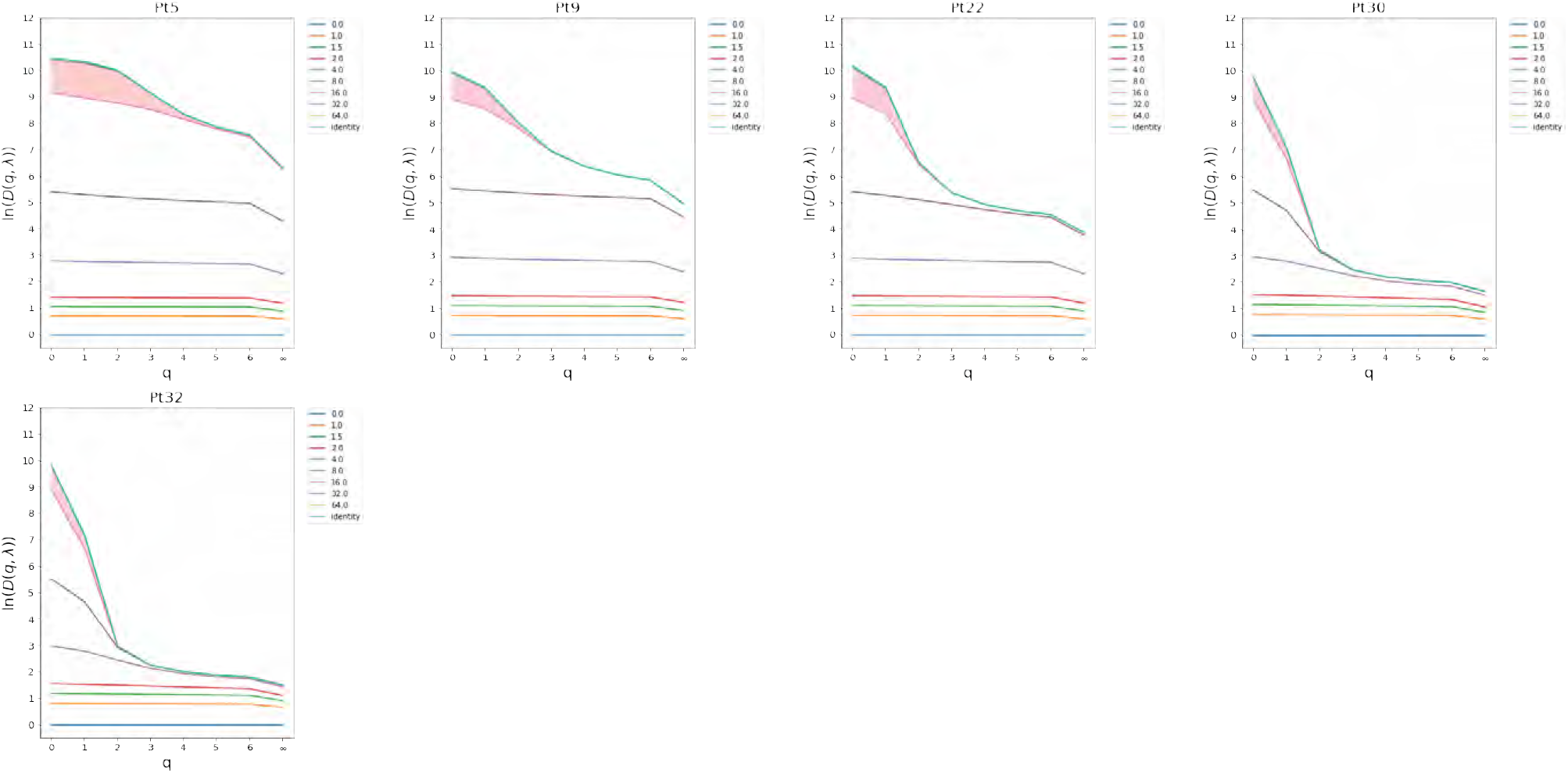
Stable Disease (SD) Day 0

**Table 12:**
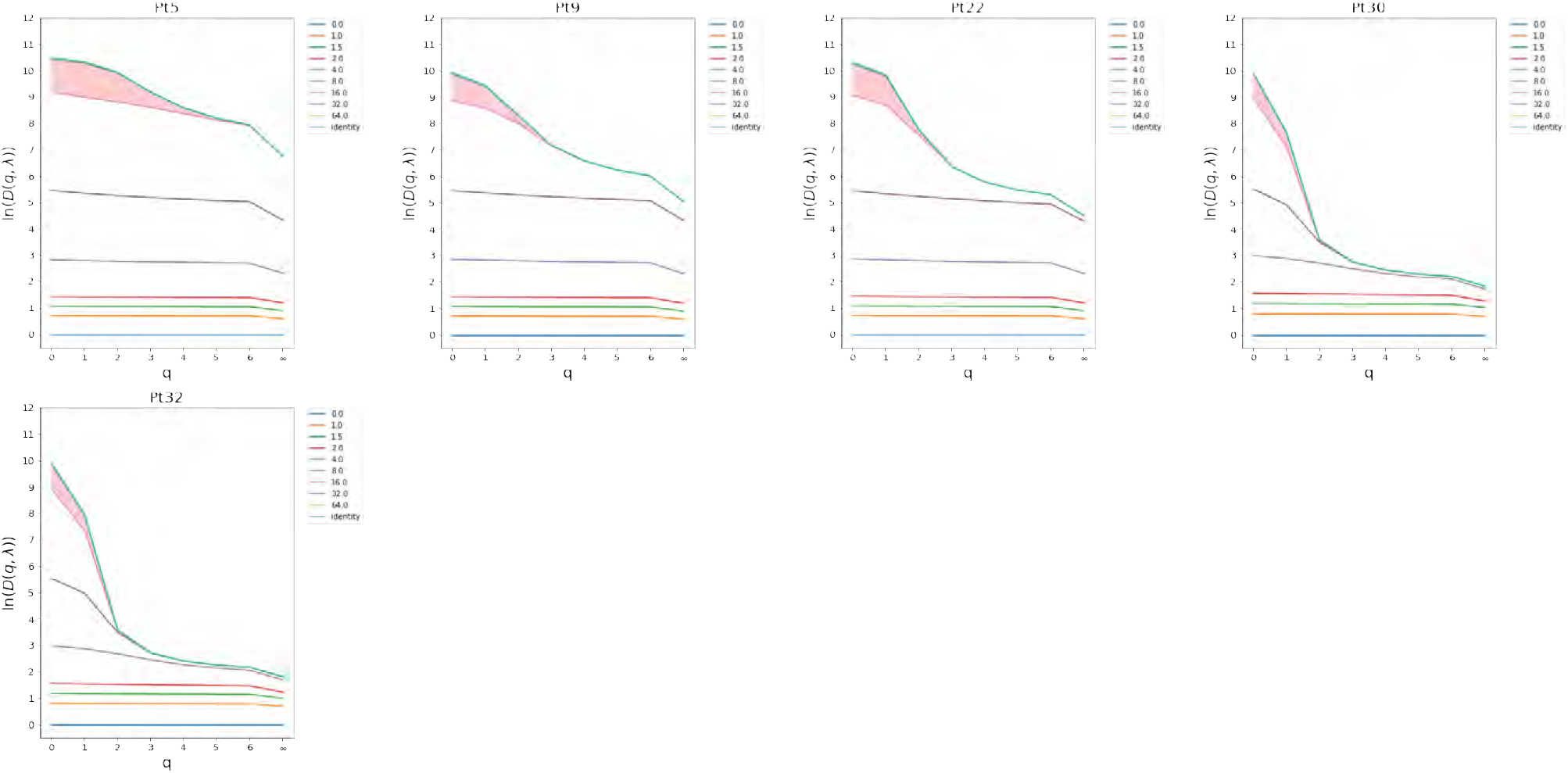
Stable Disease (SD) Day 22

**Table 13:**
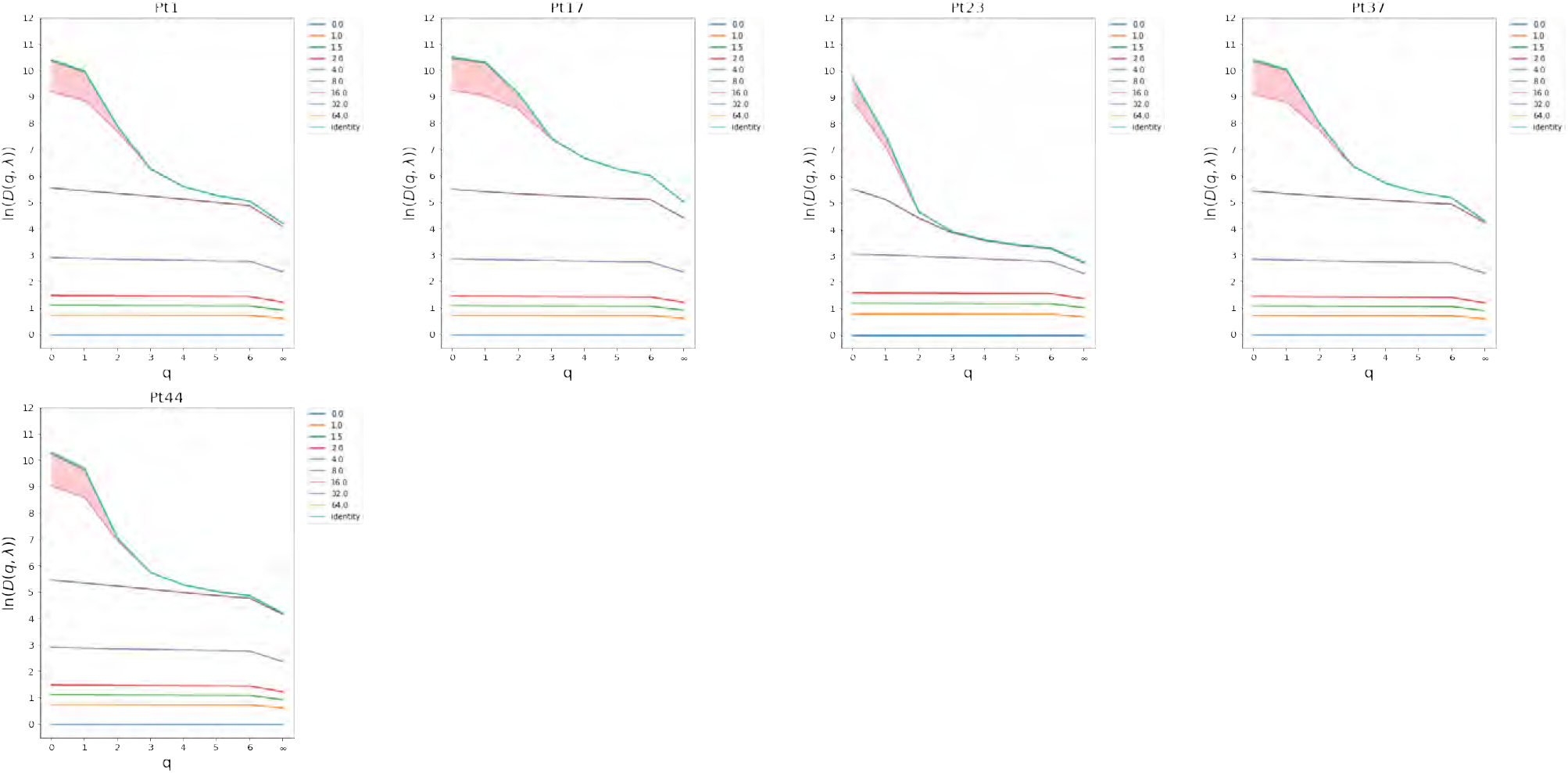
Partial Responders (PR) Day 0

**Table 14:**
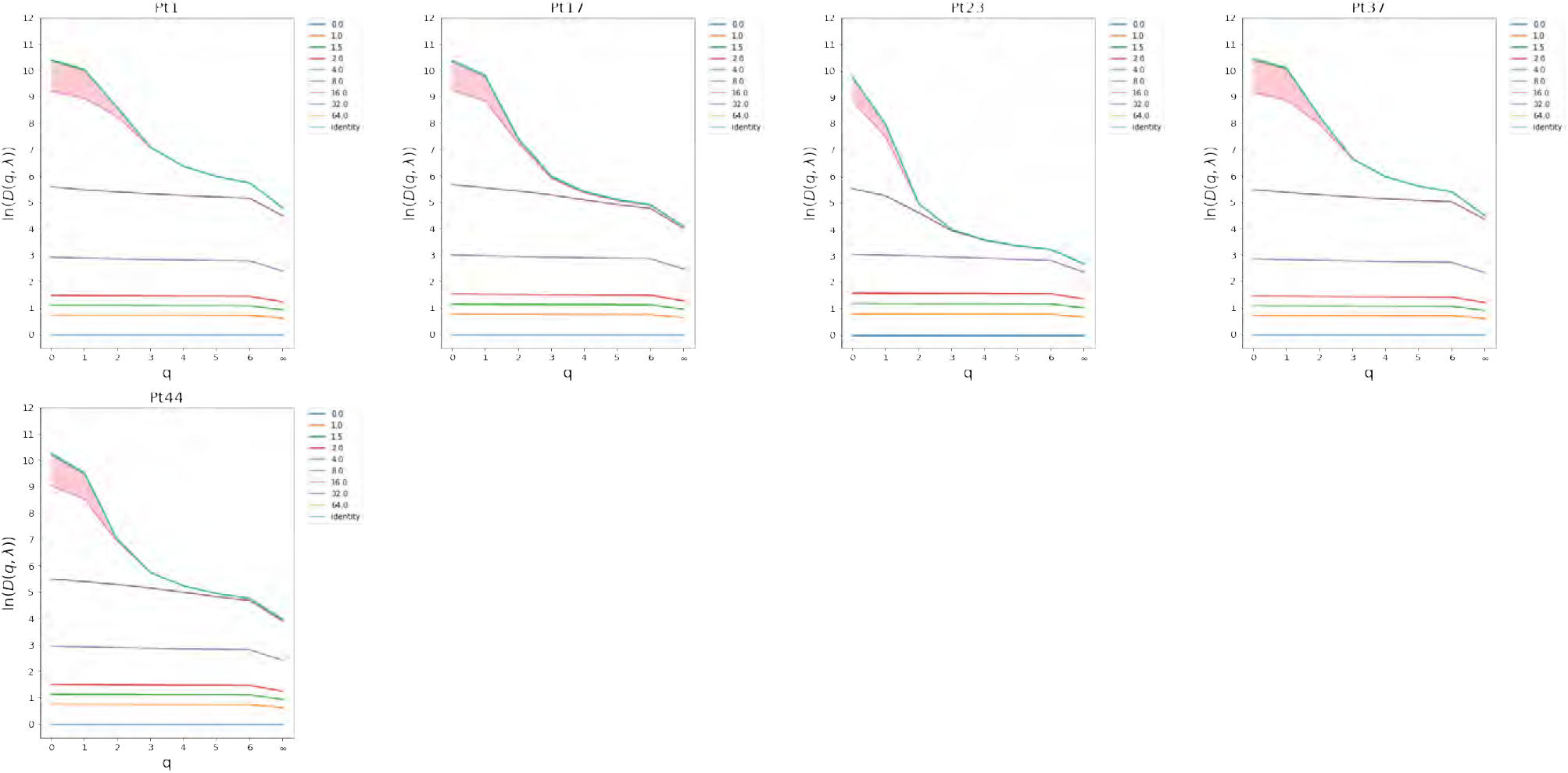
Partial Responders (PR) Day 22

**Table 15:**
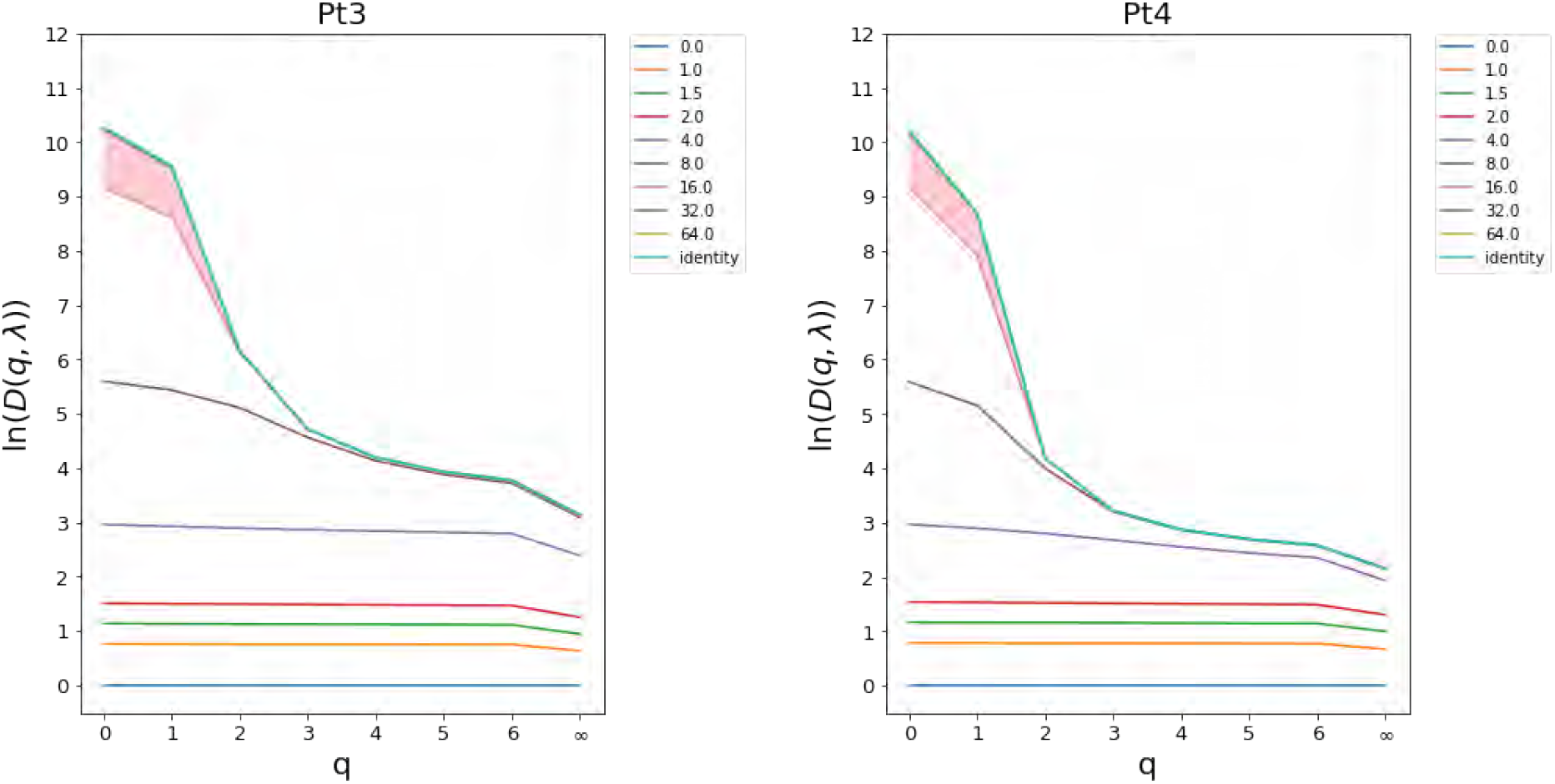
Complete Responders (CR) Day 0

**Table 16:**
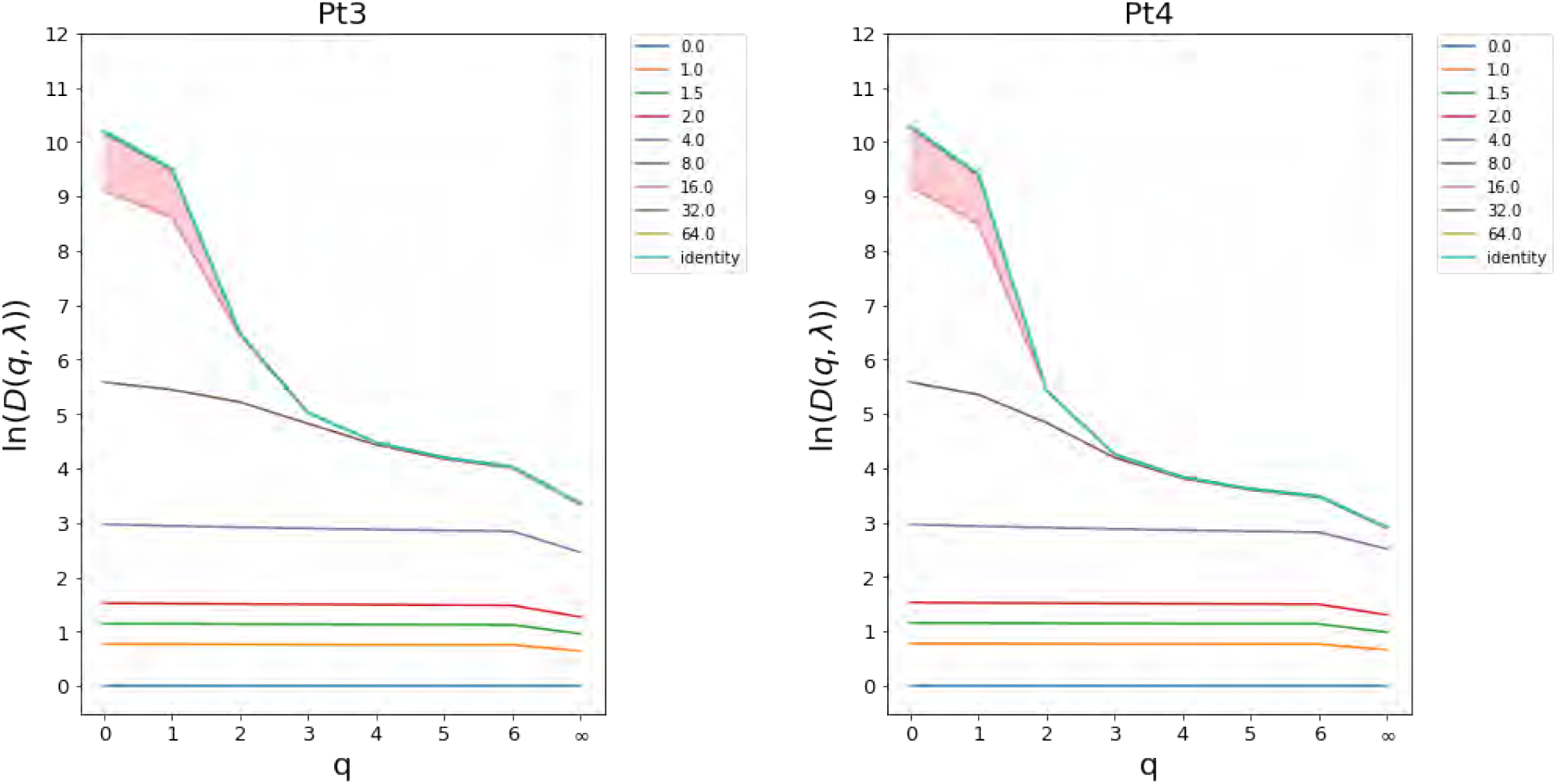
Complete Responders (CR) Day 22

#### 2.2 PCA on diversity profiles

**Figure 9:**
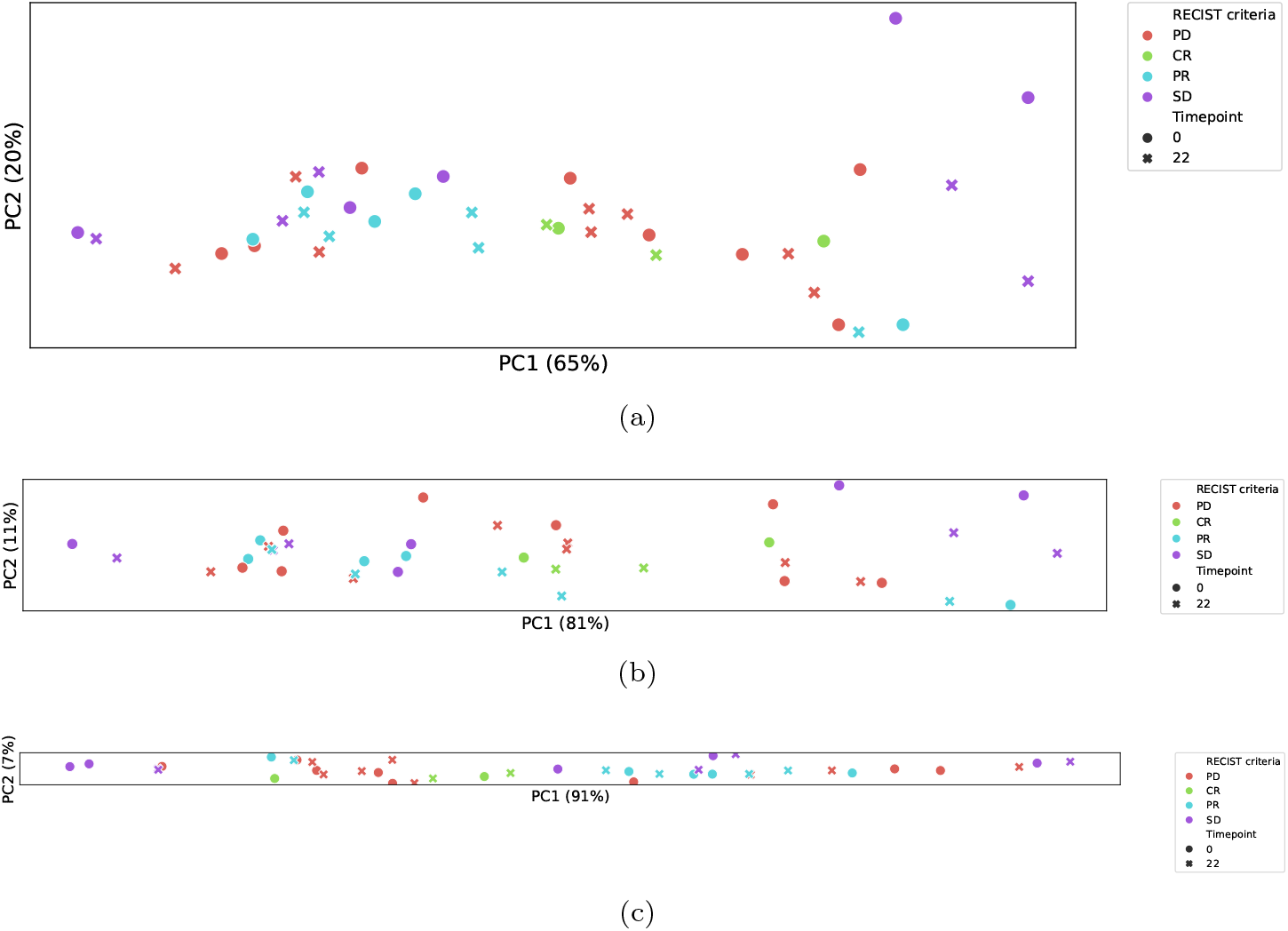
Principal Components Analysis on diversity calculated for the human dataset.The aspect ratio corresponds to variation found by PCA. **a.** PCA on features extracted from the diversity profiles constructed from the true diversity *D*(*q*, λ). **b.** PCA on values of true diversity *D*(*q*, λ). **c.** PCA on naive diversity values *D*(*q*), i.e. λ = identity.

## Notes

https://github.com/sciencisto/TCRDivER

### Competing Interest Statement

The authors have declared no competing interest.

### Summary of Updates

Author list typo.

